# The Respiratory Resistance Sensitivity Task: An Automated Method for Quantifying Respiratory Interoception and Metacognition

**DOI:** 10.1101/2021.10.14.464418

**Authors:** Niia Nikolova, Olivia Harrison, Sophie Toohey, Malthe Brændholt, Nicolas Legrand, Camile Correa, Melina Vejlø, Martin Snejbjerg Jensen, Francesca Fardo, Micah Allen

## Abstract

The ability to sense, monitor, and control respiration - e.g., respiratory interoception (henceforth, respiroception) is a core homeostatic ability. Beyond the regulation of gas exchange, enhanced awareness of respiratory sensations is directly related to psychiatric symptoms such as panic and anxiety. Indeed, chronic breathlessness (dyspnea) is associated with a fourfold increase in the risk of developing depression and anxiety, and the regulation of the breath is a key aspect of many mindfulness-based approaches to the treatment of mental illness. Physiologically speaking, the ability to accurately monitor respiratory sensations is important for optimizing cardiorespiratory function during athletic exertion, and can be a key indicator of illness. Given the important role of respiroception in mental and physical health, it is unsurprising that there is increased interest in the quantification of respiratory psychophysiology across different perceptual and metacognitive levels of the psychological hierarchy. Compared to other more popular modalities of interoception, such as in the cardiac domain, there are relatively few methods available for measuring aspects of respiroception. Existing inspiratory loading tasks are difficult to administer and frequently require expensive medical equipment, or offer poor granularity in their quantification of respiratory-related perceptual ability. To facilitate the study of respiroception, we here present a new, fully automated and computer-controlled apparatus and psychophysiological method, which can flexibly and easily measure respiratory-related interoceptive sensitivity, bias and metacognition, in as little as 30 minutes of testing, using easy to make 3D printable parts.

## Introduction

In a most general sense, perception is the detection, organisation and manipulation of sensory information. The study of internal perception, hereafter interoception, targets sensations arising from within the body, and can be defined to include perceptions originating from the visceral organs (e.g., heart, stomach) as well as processes involving them, such as respiration (Sherrington, 1952; Vaitl, 1996). Interoception influences diverse cognitive processes including decision-making, memory, and emotion (Critchley & Garfinkel, 2017; James, 1884, 1894; Nikolova et al., 2021), and is increasingly seen as a core factor in psychiatric illness (Allen, 2020; Allen et al., 2020; Khalsa et al., 2018; Khalsa & Lapidus, 2016; Owens et al., 2018). To date however, the large majority of interoception research is based within the cardiac domain, limiting our understanding of other modalities such as respiratory or gastric interoception.

Respiratory interoception (RI) is an interoceptive modality of particular interest due to its close linkage to affect (Guz, 1997; Meuret et al., 2009). Feelings of breathlessness, or dyspnea, are a core symptom of panic disorder and anxiety (Bailey, 2004; Manning & Schwartzstein, 1995; Morélot-Panzini et al., 2007), and a variety of cognitive and mindfulness-based interventions seek to improve respiratory awareness and control (Tweeddale et al., 1994). Unlike gastric or cardiac modalities, respiration is often amenable to direct conscious access, by increasing or decreasing the breathing rate, by holding the breath, or by combinations of these actions (e.g., syncopated breathing). The possibility to retrain or otherwise improve respiratory awareness and control is therefore of substantive clinical interest. Furthermore, since respiratory cycles supervene directly on both cardiac and gastric output through basic reflexes in the autonomic nervous system (an example being the cardiac sinus arrhythmia), a more thorough understanding of respiratory perception may also offer new venues for treating known interoceptive deficits in these domains (Bogaerts et al., 2008; Harrison, Marlow, et al., 2020; Harrison, Nanz, et al., 2021; Tiller et al., 1987; van Dyck et al., 2021).

To better understand RI, more precise, psychophysical measures are needed. Typically, respiratory-related interoception has been measured using resistant-detection paradigms, in which either inspiratory or expiratory breaths through a circuit can be made more or less difficult (Bennett et al., 1962; Dahme et al., 1996; Garfinkel et al., 2016; Harrison, Garfinkel, et al., 2020; Harver et al., 1993; Wiley & Zechman, 1966). For example, breathing difficulty can be manipulated by adding or subtracting static filters containing a small amount of resistance to the circuit. This technique can be paired with signal-theoretic approaches and psychophysical methods such as adaptive staircase procedures to measure perceptual sensitivity towards resistive loads, indexing the individual ability to detect pressure in the airway and diaphragm.

However, these and similar tasks suffer from several drawbacks. Previous methods for altering resistance suffer from a lack of stimulus granularity, such that the derived thresholds are extremely noisy or unreliable, and require many trials to stabilize (Harver et al., 1993). Further, resistive loads can be inherently aversive, obscuring the relationship between the objective physical detection of respiratory sensations and associated subjective affect. This combination of coarse stimulus granularity resulting in larger than necessary resistive loads, as well as the inherent aversiveness of high resistive loads, and long testing times (typically 60 + minutes in total) cumulatively make these tasks difficult for healthy participants. Crucially, these tasks are out of reach for many of the clinical populations for whom these measures would be of most importance, such as those suffering from anxiety or respiratory distress disorders (e.g., moderate to severe asthma). Additionally, classical methods only allow for the estimation of thresholds, and do not provide information regarding the slope of the psychometric function for respiratory resistance, which relates to the precision or uncertainty of the perceptual process. Clearly a more reliable, precise, and automated procedure is called for.

To achieve these aims, we designed a fully automated, 3D printable respiratory apparatus for delivering precise inspiratory and expiratory loads. Our apparatus builds on previous approaches (Garfinkel et al., 2016; Harrison, Garfinkel, et al., 2021) to enable fully computer-controlled estimation of RI thresholds, as well as improved estimation of other signal theoretic parameters such as sensitivity, bias, and metacognition. We further developed an accompanying Bayesian adaptive psychophysical approach, the respiratory resistance sensitivity task, to estimate the full respiratory psychometric function (PMF) relating stimulus level to probability of a correct response. As an initial validation of our approach, we applied this method in a sample of 32 healthy subjects. Our findings demonstrate that the respiratory resistance sensitivity task (RRST) can quickly and reliably estimate respiratory thresholds in just 20-30 minutes, with minimal subjective aversiveness. We additionally ran a direct comparison between the RRST and a recently published method (the Filter Detection Task, or FDT) in 15 healthy individuals, demonstrating much improved control in task accuracy (see Supplementary Material for details).

## Methods

### Participants

Thirty-three participants (20 females) were recruited through the Aarhus University Centre for Functionally Integrative Neuroscience Sona system participant pool. Their ages ranged from 19 to 66 years, with a mean of 28.09 years and standard deviation of 9.08 years. One participant was excluded from all data analysis due to non-convergence of the psychophysical staircases, resulting in 32 participants for the final analyses. For the analysis of type 2 performance, a further two participants were excluded due to too many extreme confidence ratings. All participants had normal or corrected-to-normal vision, and fluent English or Danish proficiency. Furthermore, participants did not have a current psychiatric diagnosis, and did not use drugs or medications other than hormonal contraceptives or over the counter antihistamines. In total, the experiment lasted 60-75 minutes, and participants received 350 DKK in compensation for taking part. Since data collection was performed during the COVID-19 pandemic (January - March 2021), participants were required to present a negative COVID-19 test made within 48 hours of the day of the experiment. The experimenter wore a face mask and/or shield at all times, and participants wore a face mask when not performing the task. The study was approved by the local Region Midtjylland Ethics Committee and was carried out in accordance with the Declaration of Helsinki.

### Device specification

To enable the precise, automated delivery of respiratory resistive loads, we developed a novel apparatus based on previous resistive load tasks (Garfinkel et al., 2016; Harrison, Garfinkel, et al., 2020). The primary mechanism of the device is a 3D-printed housing which secures a section of flexible tubing against a wedge. This tubing is then connected to a sanitary, hospital grade respiratory circuit by custom-fit couplers. This wedge is then connected by a screw to a step motor, and is then driven forwards or backwards against the flexible tubing by the motor. The motor itself is connected to an Arduino circuit board which is programmed with the instructions for converting digital inputs into discrete steps along the screw. Thus, by delivering electronic commands to the step motor, the wedge moves forwards or backwards against the length of tubing, resulting in a reliable stepwise compression or relaxation, i.e., an increase or decrease of the static resistance through the full respiratory circuit. At the participant end of the device a Hans Rudolph 2-way non-rebreathing t-valve connects the respiratory tubing to a PowerBreathe TrySafe filtered mouthpiece, through which the participant breathes.

A stepper motor is controlled by a TMC driver and Teensy 3.1 microcontroller. The motor is securely attached to a threaded TR8×2 leadscrew by a coupler. The TR8×2 leadscrew and nut are used due to their design for use in 3D printers, and are therefore made to withstand repetitive and continuous movement. A motor damper and rubber feet are further used to reduce vibration and noise from the device.

The RRST software was written using MatLab (2020a), using PsychToolBox (Brainard, 1997; Kleiner et al., 2007; Pelli, 1997) and the Palamedes toolbox (Prins & Kingdom, 2018). All task code, associated data analysis scripts, and anonymized participant data are publically available on our GitHub page: https://github.com/embodied-computation-group/RespiroceptionMethodsPaper

### Set-up

Participants performed the RRST while seated in front of a computer at a height-adjustable desk. The user-end of the breathing circuit with the single use mouthpiece was positioned and held in place in front of the participant using a desktop microphone stand. Participants were encouraged to adjust the height of the desk so as to be able to comfortably lean forward and inhale through the mouthpiece while performing the task. Several precautions were taken to eliminate non-respiratory cues, such as visual, auditory and tactile cues associated with the movement of the device. First, participants were fitted with over-ear headphones playing continuous ‘rain’ noise selected to mask sounds generated by the device. Second, the compressed end of the breathing circuit and the device were placed within a box located to the side of the participant, such that they were not able to see any movement. Third, to eliminate vibrations caused by the stepper motor, it was fitted with a damper and rubber feet were attached to the device. The apparatus was further placed on top of thick foam padding inside the box, which was positioned on a surface detached from the participants’ desk. Finally, the position of the load was changed to a random value during each inter-trial interval to decorrelate the duration of movement to the stimulus intensity on the subsequent trial.

At the start of the first session, participants completed a tutorial session. This introduced the trial structure in a stepwise manner, using two trials with accuracy feedback, and six additional trials of varying difficulty without feedback. This ensured consistency in the task instructions, and gave participants a chance to learn and adapt to the trial structure and breathing pace before the start of the adaptive staircases. The tutorial lasted about 5 minutes, the QUEST staircase 15 minutes, and the Psi staircase 30 minutes including breaks.

#### Trial structure

To measure respiratory resistance sensitivity, we used a two-interval forced-choice (i.e., temporal two-alternative forced-choice) design (2IFC). When interested in determining measures of perceptual sensitivity, a 2IFC design is preferable to one where a single stimulus is presented on each trial because they are less susceptible to biases in the decision process. Indeed, it has been shown that participants can voluntarily shift their psychometric curve on a single-interval forced-choice task, without altering their sensitivity (Morgan et al., 2012).

On each trial, participants took two breaths, where one (the stimulus) had a resistive load applied to the breathing circuit while the other (the standard) did not. Which of the two breaths contained the resistive load was determined pseudorandomly. After taking the two consecutive inhalations, participants decided whether the first or the second breath was more difficult, and entered their response using the left and right buttons of a computer mouse. They were then asked to rate how confident they were in their decision by using the mouse to move a slider on a visual analogue scale (VAS) ranging from ‘Guess’ to ‘Certain’ and confirming the answer by a left-click on the mouse. The maximum response time for the decision was 3 seconds, and 5 seconds for the confidence rating. If no response was made during this time, the trial was repeated.

The inhalation pace was cued by a visual stimulus: the outline of a circle appeared on the screen for 200ms, then an expanding gaussian ring was displayed for 800ms, during which time the ring grew to occupy the space within the circle. Participants were instructed to pace their breaths to the expansion of the ring, to breathe “sharply and shallowly” and that inhales should last just under a second. We recommended participants to keep their mouth on the mouthpiece for the duration of each trial, exhaling through the nose between the two intervals. This instruction was provided to minimize rebreathing of air from within the circuit. We did not control the breathing strategy used by participants further (e.g., participants were free to exhale through the breathing circuit or outside it). To avoid hyperventilation, the task contained forced breaks of at least 2 minutes in duration every 20 trials (ca. every 5 minutes of testing). Participants were instructed to use these breaks to stand up, move around and breathe naturally. Prior to each break, participants rated how aversive they found the task stimuli using a VAS from ‘Not unpleasant’ to ‘Very unpleasant’. At the end of the session, participants responded to subjective experience questions assessing ‘dizziness/light-headedness’, ‘breathlessness’, and ‘presence of asthma symptoms’.

### Psychophysical methods

To estimate participants’ sensitivity to detecting small increments of obstruction of airflow on inhalations, we used established psychophysical methods to measure the PMF threshold and slope. The threshold corresponds to the Weibull parameter *α*, denoting the stimulus value at which the probability of responding correctly is 75%. The slope corresponds to the Weibull parameter *β*, approximating the signal uncertainty. *β* is proportional to the gradient of the function at stimulus x = *α* when *Ψ* is plotted on log units. Varying *α* produces a left - or rightward shift of the PMF, while varying *β* changes its steepness, with larger *β* values resulting in a steeper function. In the context of the RRST, the threshold corresponds to the detection sensitivity, while the slope gives information about their uncertainty, or the response variance with changes in stimulus intensity.

The Weibull function is often modified to include a lapse parameter *λ* so that the function asymptotes at a performance level of (1 - *λ*), allowing for lapses due to inattention or motor error. The Weibull function *Ψ_W_* relating the predicted proportion correct *P(correct)_W_*, is then given by:

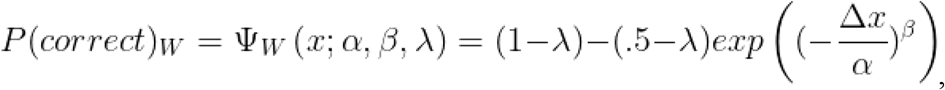

Where *x* represents a value on the stimulus dimension, *α* denotes the threshold, *β* the slope, and *λ* the lapse rate.

Each participant completed two sessions of the RRST, using different psychometric methods with counterbalanced order to evaluate the internal reliability of the task, namely the QUEST staircase (Watson & Pelli, 1983) and Psi (Kontsevich & Tyler, 1999). Next, we describe these two methods, and our use of them for the RRST. Both QUEST and Psi are Bayesian adaptive psychophysical methods, and use prior information provided by the experimenter (from previous experiments, or from literature) in addition to information from all preceding trials in order to guide the placement of the next stimulus level in order to efficiently obtain estimates of the PMF parameters of interest. Whereas QUEST estimates the threshold only, Psi can be used to estimate both threshold and slope. The staircase procedures were initiated to run in units of wedge displacement (i.e., proportional to units of motor rotation) ranging between 0 and 18 mm, and these units were also used for all analyses. To aid interpretability, we transform these values into percentage obstruction for the figures. In this study, QUEST was initiated with a uniform prior distribution (range 0 to 18mm) over the threshold. Two interleaved QUEST staircases of 30 trials each were run within a single (60 trial long) session, and the mean of the posterior distribution was used as the threshold estimate for each staircase (King-Smith et al., 1994). Subsequently, the mean of the two interleaved staircases was taken as the QUEST threshold.

For every trial, Psi considers the range of possible stimulus intensities to present. For each intensity, Psi computes the probability of a correct versus incorrect response as well as the expected entropy resulting from either response. The stimulus level for which the expected entropy is lowest is then presented on each trial. Following the observer’s response, Psi then uses the posterior distribution to recalculate the PMF that best fits the data from all previous trials. Here, Psi was initiated with the following prior parameters: Weibull PMF, *α* uniform 0 to 18, *β* uniform log(1) to log(16), guess rate *γ*=0.5, and lapse rate *λ*=0.02. The guess and lapse rate parameters are fixed and determine the lower and upper asymptotes, respectively. Individual and group PMFs relating stimulus level (% obstruction) to the probability of a correct response were fit using a Bayesian criterion. The search grid was defined with the same parameters used for the priors for the Psi staircase, with wide distributions for the threshold and slope, and fixed values for the guess and lapse rates.

### Physiological measures

To determine the relationship between the degree of tube obstruction and effective static resistance produced in the airway, it is important to obtain measures of the mechanical properties of the air circuit. To this end, we ran the task at a second site (University of Otago, New Zealand), enabling us to record physiological measures alongside the RRST (see Supplementary Material - Task validation using physiological measures for further details). We measured airflow and differential pressure at a constant airflow of 0.95 L/s, and used these to calculate the resistance at each level of obstruction. Resistance corresponds to the force opposing the flow of air through the circuit, and remains constant for a given degree of compression on the circuit. The force of inspiration can however change the differential pressure exerted across the area of compression, leading to changes in inspiratory flow. The relationship between resistance, pressure and flow is given by Ohm’s law

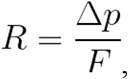

where resistance R is determined by pressure Δp and flow rate F. Measures of differential pressure, and airflow were collected both for human participants performing the task and using a constant flow device. Physiological measures of mouth pressure and inspiratory flow were measured using an ADInstruments pressure gauge coupled to a bridge amplifier, and inspiratory flow was measured using an ADInstruments spirometer with a 300 L flow head. Physiological traces were recorded via a 16-channel PowerLab (ADInstruments, New Zealand), and analyzed within LabChart (version 8; ADInstruments).

The constant airflow was generated using a custom-built device that blew a steady stream of room-temperature air through the breathing circuit. Since the effective pressure depends on the characteristics (e.g., depth, strength, acceleration) of each inhale, the constant flow data was used to analyze the resistance generated at different levels of obstruction without breath-to-breath variability. This also allowed us to visualize the variance and drift in resistance across trials. The constant flow device was attached to the circuit instead of the mouthpiece section. LabChart software was used to record the pressure and flow data while the RRST device moved in increments between positions corresponding to 0 and 100% obstruction. Each obstruction measure was interspersed with a 0% obstruction position, such that the device moved in a similar manner to what is employed during experimental tests. Therefore, each of 17 obstruction steps were interleaved with the device returning to 0% obstruction, and the process was repeated three times in total.

### Analysis

Statistical analyses and fitting of PMFs were performed using MatLab (2020a) and the Palamedes toolbox (Prins & Kingdom, 2018). We used Pearson correlation to evaluate the correlations between type 1 and type 2 variables, and to determine between-method agreement. The effect of trial accuracy on response times and stimulus level was tested using t-tests assuming unequal variances (F-test for equality of variance). Exploratory correlations between threshold and mean aversiveness judgements and subjective ratings of dizziness, breathlessness were tested using Spearman correlation. The figures were created using MatLab, and the distributions for threshold, slope, response times and stimulus levels were made using raincloud plots (Allen, Poggiali, et al., 2019).

We used a signal theoretic approach to evaluate perceptual and metacognitive sensitivity, to reduce the influence of response biases and estimate metacognitive performance independently from perceptual sensitivity. For perceptual (type 1) performance, trials on which the presence of a resistance was identified were coded as correct, for example if the first alternative was chosen, given that the resistance load was indeed applied on the first breath. Conversely, incorrect trials were those on which the standard trial (without resistance load) was chosen. For metacognitive (type 2) performance, the confidence ratings were first converted from raw ratings of 0 to 100 into 4 equally spaced bins. Then metacognitive signal theoretic measures can be defined; ‘hits’ are marked as trials on which the type 1 response was correct, and the confidence rating was high, whereas ‘misses’ are trials on which the type 1 response was incorrect but the confidence was high (Fleming & Lau, 2014; Maniscalco & Lau, 2012). In this way the area under the type 2 receiver operating characteristic (aROC) curve can be determined, which estimates the metacognitive sensitivity while accounting for a bias to over or under confidence. The type 1 and 2 parameters were calculated using the HMeta-d toolbox (Fleming, 2017). We used the area under the type 2 receiver operating characteristic (aROC) as the main metacognitive measure here because performance was tightly controlled at around 80% by the Psi staircase (Fleming & Lau, 2014; Maniscalco & Lau, 2012).

For analysis of the physiological measures recorded during constant airflow application, pressure and flow measures were averaged across a 2 second interval at each level of obstruction for each of three experimental runs. Resistance was calculated by dividing the change in pressure by the average flow at each obstruction step. Percentage obstruction was then plotted against the measured resistance values. Additionally, as an exponential relationship was observed between percentage obstruction and resistance, the resistance values were log-transformed and then re-plotted against percentage obstruction, where a linear relationship could then be quantified.

## Results

### Psychometric results

We first evaluated the staircase convergence and threshold estimates for the QUEST and Psi staircase methods. As similar results were observed from the two methods, the following results are presented for the Psi staircase session only (see type 1 performance - Performance for Psi and QUEST staircases for details). Individual and group PMFs relating stimulus level (% obstruction) to probability of a correct response were fit using a Bayesian criterion. The search grid was defined with the same parameters used for the priors for the Psi staircase, with wide distributions for the threshold and slope, and fixed values for the guess and lapse rates (see Methods, Psychophysical methods above). The mean group threshold (*α*) was 65.61% obstruction (SD = 8.47%), and the individual threshold values ranged from 56.71 to 89.79%, as shown in **Figure 2 C & D**. Slope estimates also exhibited large inter-subject variability, with a mean of 7.80, standard deviation of 2.61, and range from 3.12 to 12.37.

**Figure 1 Legend:**
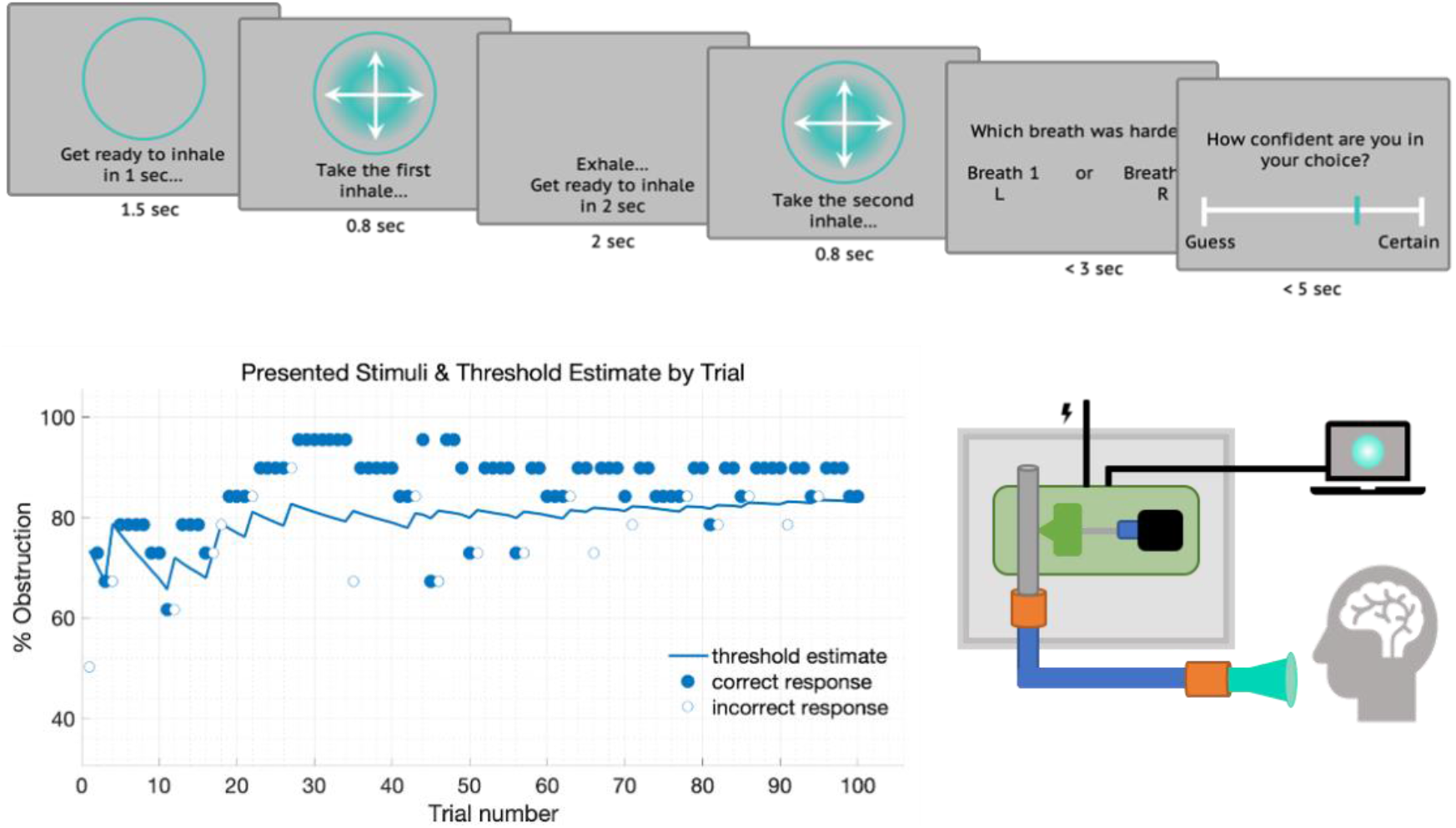
The Respiratory Resistance Discrimination Task. A) Trial schematic depicting the 2-interval forced choice (2IFC) design of the task. On each trial participants view a circular cue instructing them to prepare to inhale. The circle then blinks and begins expanding, with the participant instructed to sharply inhale with the expansion of the circle. The participant then exhales, and a second similarly guided breath is conducted. This procedure of pacing the participant’s breathing via visual cues is a novel feature of the RRST, and is intended to reduce intra- and inter-subject variance in respiratory effort. Following the two breaths, the participant indicates by keyboard press whether the first or second breath was more difficult. B) Sample single subject data, illustrating the psychophysical procedure. On each trial, one of two breaths is randomly signal minus (s-), such that the compression wedge is at resting baseline (0% obstruction) with no added resistance, and the other signal plus (s+) with some level of compression determined by the staircase procedure. The procedure rapidly hones in on a threshold estimate using a Bayesian procedure (psi); in this example the participant threshold of approximately 80% obstruction is found within just 20 trials. C) Schematic illustrating the design of the automated resistive load apparatus (see Supplementary Material 1 - Detailed schematic for details).

**Figure 2:**
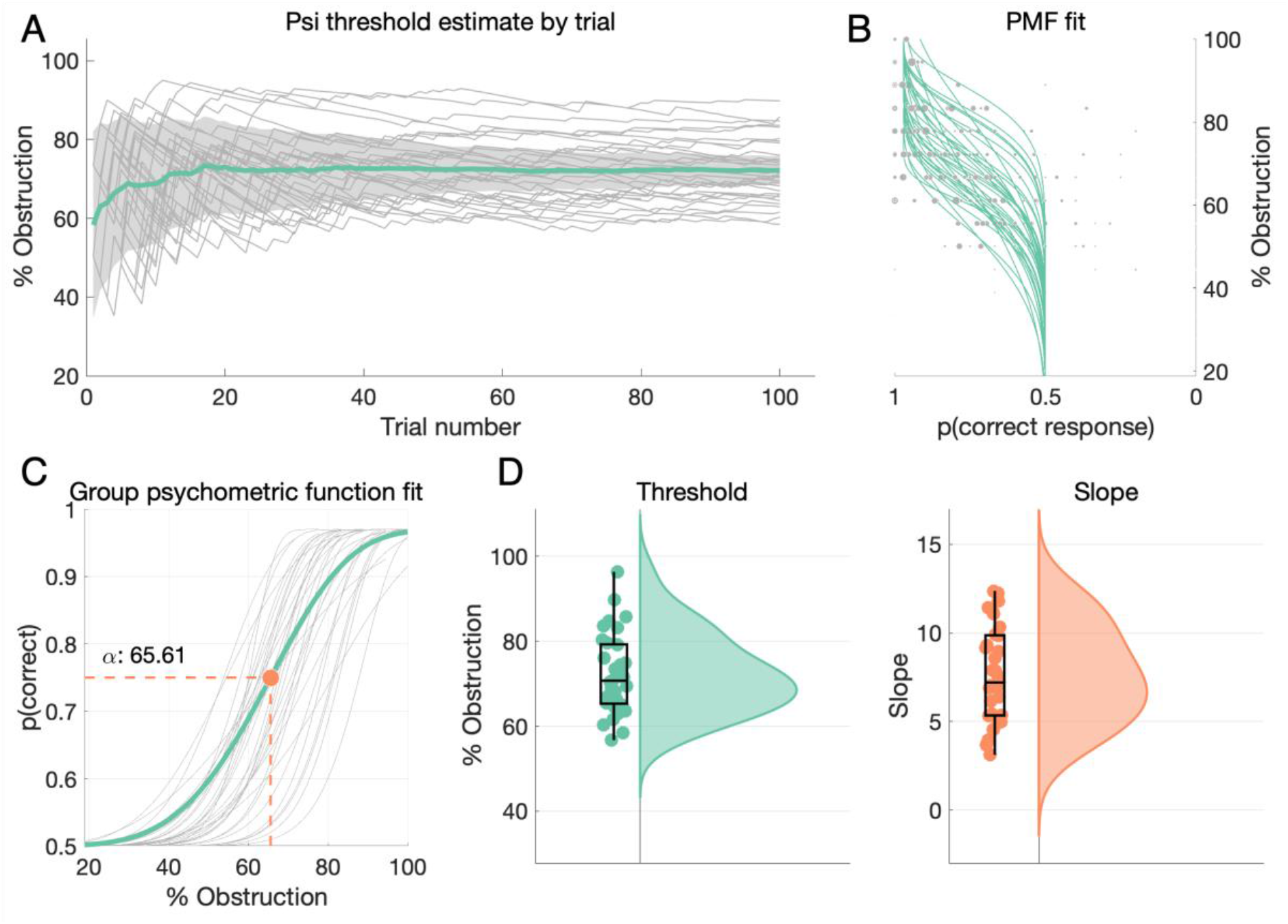
Psychometric task results. **A)** Plot depicting trial-by-trial Psi threshold estimates for all participants. Light gray lines depict individual stimulus traces indicating the % of tube obstruction on the stimulus breath for each trial, the thick green line represents grand mean stimulus on each trial +- SEM. In general, threshold estimates stabilize around trials 20-40 for all participants. **B)** PMF fits for all subjects. The green lines depict individuals’ PMF fits, and grey points show stimulus levels presented, where the dot size indicates the number of times presented. **C)** Grand mean psychometric fit (green) overlaid on individual PMF fits (grey), demonstrating that average respiratory thresholds are around 66% airway obstruction, with substantive inter-individual variance around this value. **D)** Raincloud plots (Allen, Poggiali, et al., 2019) depicting individual threshold (green) and slope (orange) estimates for all subjects.

### Type 1 performance - perception

The RRST method estimates subject thresholds by presenting stimuli at various points around the estimated decision function. As a partial validation of this procedure, we first examined the relationship between choice accuracy, stimulus intensity, and reaction time. If estimated thresholds are reasonably well estimated, then we would expect responses to be significantly faster for correct versus error trials, and to observe that correct versus incorrect trials are generally associated with higher stimulus intensity levels. Indeed, when examining the effect of accuracy on reaction times (RT) for Psi staircase sessions, we observe that RTs are significantly lower for correct versus incorrect trials, *t*(42.25) = −3.34, p < 0.01 (Figure 3). Stimulus intensity levels on correct trials were significantly greater than on incorrect trials, *t*(60) = 4.66, p < 0.01 (Figure 3). These results indicate that near-threshold signal-present stimuli (i.e., resistances just above and below a participants’ threshold) modulate both processing time and response accuracy as expected.

Similar results are obtained for the Quest staircase, with lower RTs for correct compared to incorrect trials *t*(49.8) = −3.35, p < 0.01, and higher stimulus intensities on correct versus incorrect trials, *t*(60) = 3.82, p < 0.01.

**Figure 3:**
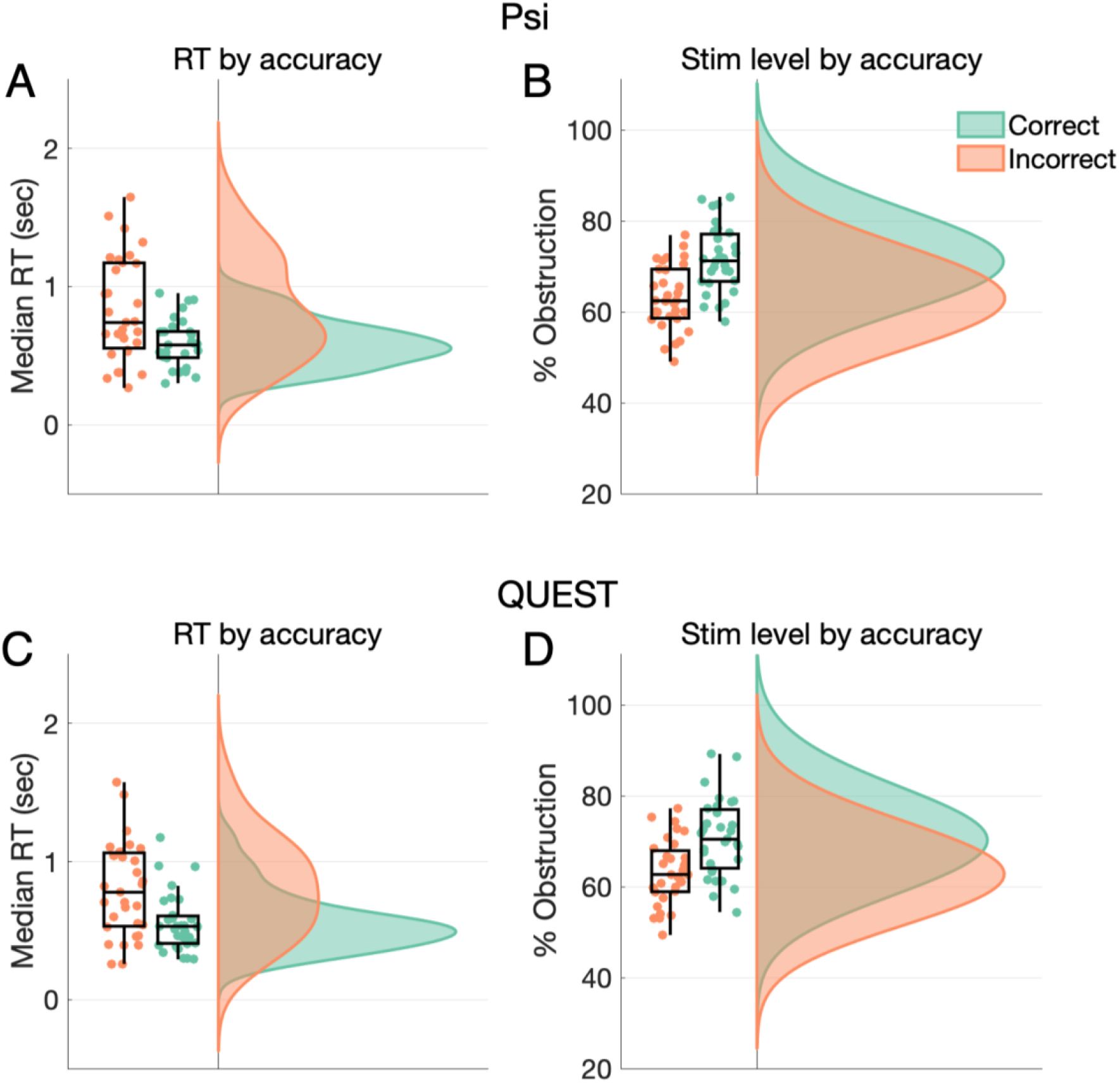
Type 1 performance on Psi & QUEST methods. Raincloud plots of reaction times (RT, left panel) and stimulus level (right panel) by accuracy (correct vs. incorrect) for the Psi (upper panel) and QUEST (lower panel) staircase methods. **A) & C)** Median RTs presented for each subject, for correct (green) and incorrect (orange) trials showing that RTs on correct trials are lower than on incorrect trials. **B) & D)** Average stimulus levels presented for each subject, for correct and incorrect trials, showing that stimuli were higher (i.e., easier) on correct trials. These results indicate good overall convergence of estimated psychometric thresholds.

### Task reliability

To determine the number of trials needed for staircase convergence, we analyzed measures of how far-removed current staircase estimates were from the final threshold estimate at the end of the session. On each trial, the Psi method determines a standard error (SE) for each PMF parameter it is estimating. The progression of SE estimates for the threshold and slope over trials are shown in **Figure 4 A & B**. Visual inspection shows that threshold SEs shrink the most up to around trial 50, whereas slope SEs show a decreasing trend even at trial 100. This suggests that over 100 trials may be necessary to accurately estimate the slope parameter of the PMF. For QUEST, we calculated the difference between the stimulus (% obstruction) level presented on each trial and the final threshold estimate (see **Figure 4 C**). This measure asymptotes around trial 20, however care should be taken in the interpretation as this measure is not directly comparable to the SE estimates from Psi above.

**Figure 4:**
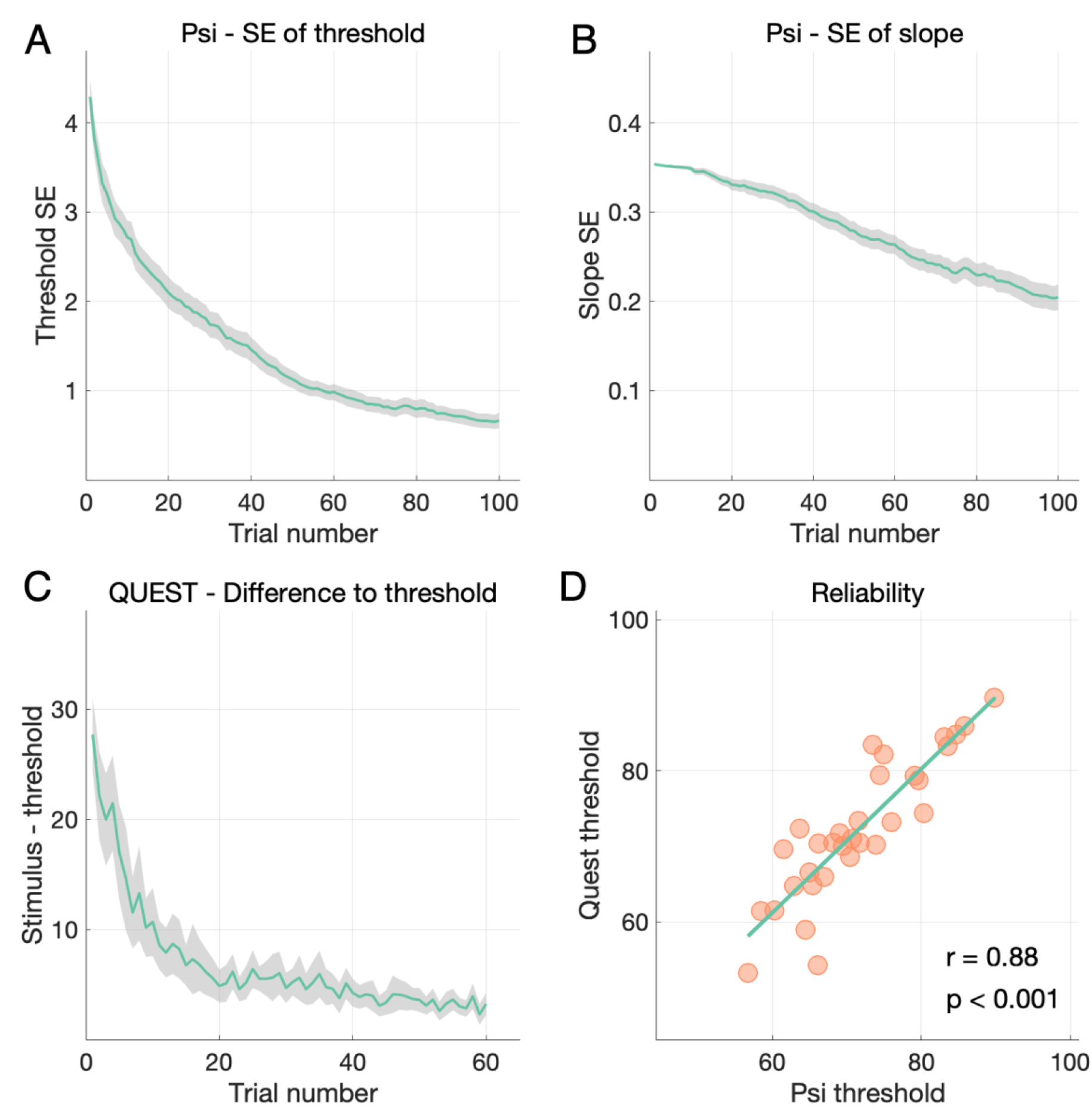
Staircase convergence and Task Reliability. **A)** Standard errors of the threshold estimate by trial number obtained with Psi indicate that reliable threshold estimates are derived within 20-50 trials. **B)** Standard errors of the slope estimate by trial number show that slope uncertainty drops linearly as a function of trials, indicating that slope estimates may benefit from higher trial numbers and/or hierarchical modelling. **C)** The difference to the final threshold estimates by trial number obtained with QUEST indicates that threshold estimates also converge within 20-50 trials. **D)** Correlation plot across the two (counter-balanced) QUEST and Psi threshold estimates indicates a high within-subject reliability of respiroceptive thresholds, regardless of estimation technique.

The within-subject reliability of the thresholds obtained using the RRST was assessed by a Pearson correlation on the estimates obtained from the Psi and QUEST staircase methods. Thresholds on the two interleaved sessions were found to be strongly correlated, r(31) = 0.88, p < 0.001 indicating high consistency between the two methods.

### Type 2 performance - metacognition

To assess respiroceptive metacognition, we asked participants to rate their confidence in their decision on every trial. We here describe the results of the metacognition analysis, based only on trials from the Psi session only. Confidence ratings were generally higher for easier trials (i.e., trials on which the obstruction was greater, see **Figure 5 A**), and there was a dissociation between ratings on correct and incorrect trials, such that confidence was lower on incorrect trials and higher on correct trials (see **Figure 5 B**). To estimate individuals’ metacognitive ability, we calculated the area under the type 2 ROC curves (see **Figure 5 C**, the aROC is the area between the identity diagonal and the ROC curve), which corresponds to the ability to associate confidence to perceptual performance. While the Psi staircase held perceptual performance constant at a level of 76 - 84% task accuracy, metacognitive ability (aROC) varied substantially, ranging from 0.48 - 0.85 (mean = 0.70, SD = 0.09, see **Figure 5 D**). Furthermore, the type 1 and type 2 performance measures were not correlated (r(31) = −0.03, p = 0.86). These results highlight the unique ability of the RRST approach to titrate respiroceptive performance and thus enhance the specific estimate of interoceptive metacognitive ability.

**Figure 5:**
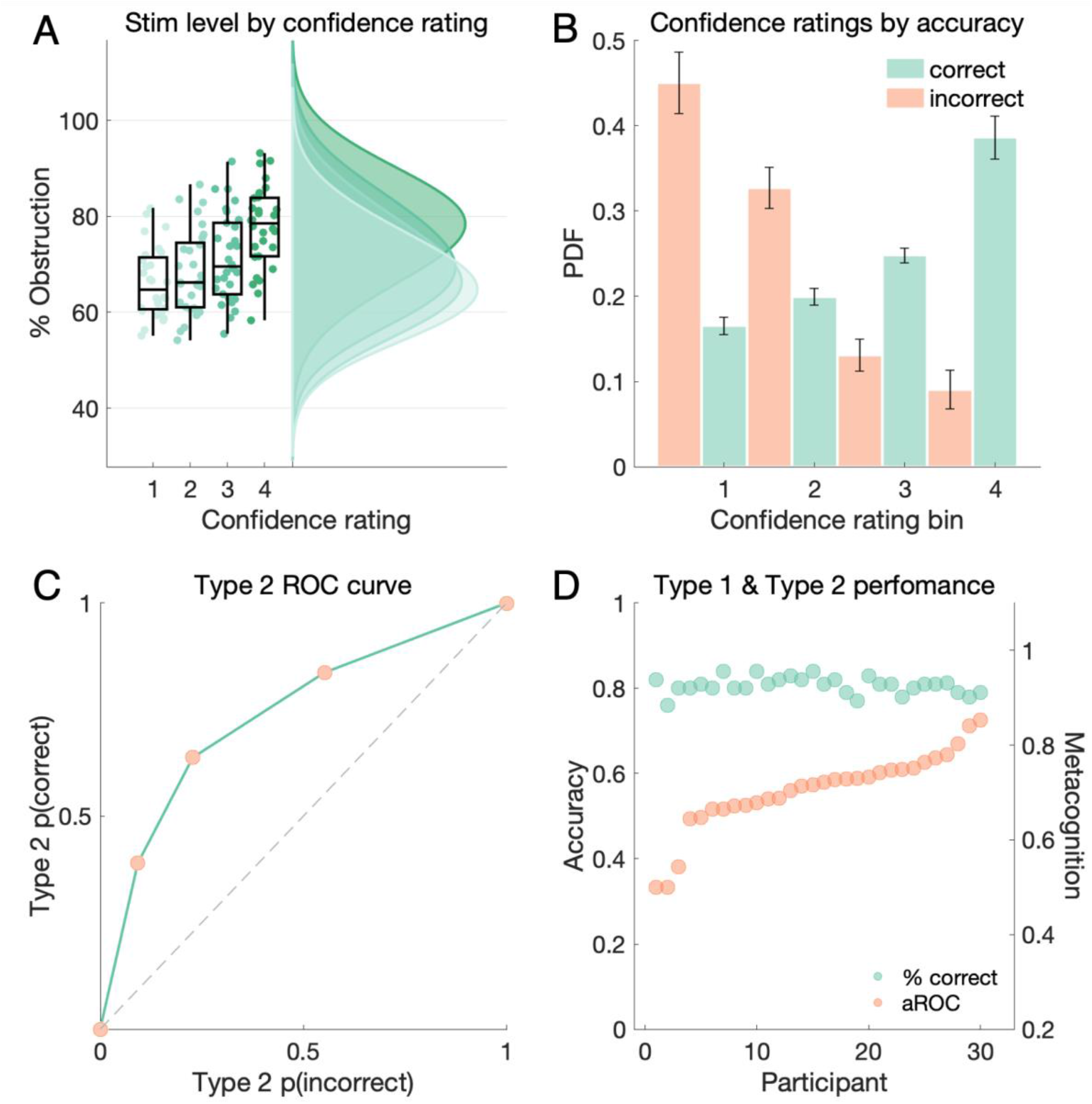
Type 2 performance. **A)** Raincloud plot showing stimulus level by confidence rating, showing higher confidence ratings for stimuli with greater resistance. **B)** Histogram of binned confidence ratings for correct (green) and incorrect (orange) trials. Generally, participants showed high metacognitive sensitivity, as seen in the dissociation between the correct and incorrect trial ratings. **C)** Type 2 ROC curve, averaged over participants, showing good respiroceptive type 2 performance. **D)** Type 1 (accuracy, green) and type 2 (aROC, orange) performance, sorted by each participant’s aROC. Participants show substantial variations in metacognition while type 1 accuracy is held relatively constant by the Psi staircase procedure.

**Figure 6:**
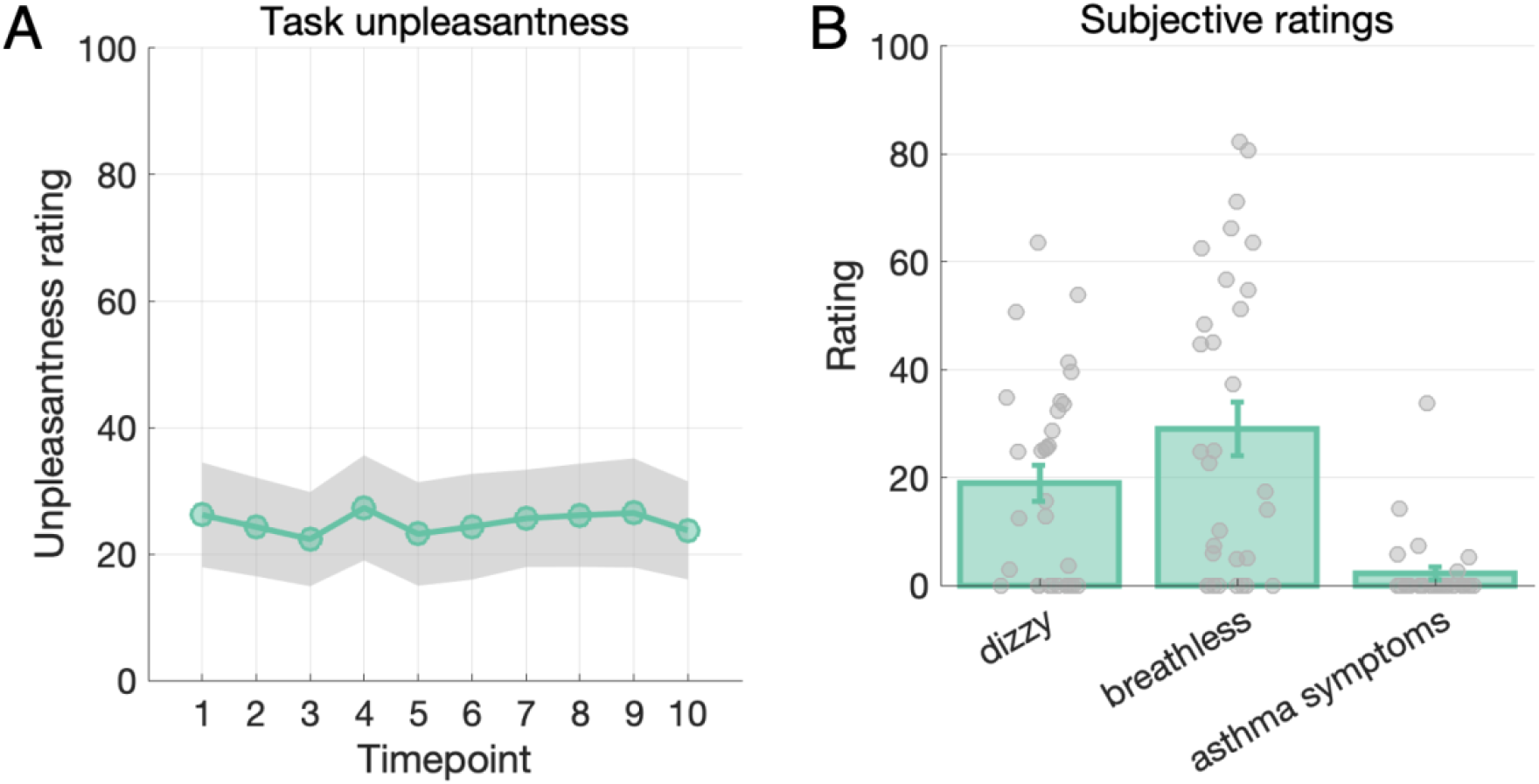
Task tolerance & subjective ratings. **A)** Plot depicting aversiveness ratings across all 10 blocks (timepoints) of testing. Mean aversiveness ratings after each block are shown in green, and the shaded gray area represents standard error of the mean (SEM). Each block comprised 20 trials, for a total testing time of approximately 45 minutes. Participants on average reported roughly 20% stimulus aversiveness (out of 100 total), which remains stable throughout the testing period. This indicates that the stimuli were mildly unpleasant and that extended testing time did not increase task adversity within these limits. **B)** Plot depicting mean dizziness, breathlessness and asthma symptoms across participants. Bar height represents mean ratings, error bars denote SEM, and gray circles show individual participants’ ratings. In general, participants showed low levels of these adverse effects following 1 full hour of testing, indicating good tolerability of the task.

### Task tolerance

We wanted to assess whether participants found the task unpleasant or aversive, and whether this changed over the course of the testing session. **Figure 5 A** shows the aversiveness ratings over 6 blocks, or 45 minutes, of testing. Mean displeasure ratings were around 25% (SD = 1.61), indicating that the task was on average mildly aversive and that this remained constant throughout the testing session. Average dizziness/light-headedness, breathlessness, and severity of asthma symptoms are shown in **Figure 5B**, along with data points indicating individual ratings. Ratings of aversiveness were found to correlate strongly with those of dizziness, but not with other subjective ratings, or with perceptual or metacognitive task parameters (see **Supplementary Figure 2** for details).

### Inspiratory resistance

In **Figure 7**, we show the measured resistance over levels of obstruction. The relationship between obstruction and resistance is exponential, with resistance increasing dramatically from about 70% obstruction (**Figure 7A**). The relationship between the natural log of resistance and obstruction is linear (**Figure 7B**, slope = 0.44, p < 0.0001, R^2^= 0.88), with some variability at extremely low circuit obstruction and deviation from linearity at extremely high circuit obstruction. Variability at low circuit obstruction levels is likely due to measurement-associated error at very low levels of resistance.

**Figure 7:**
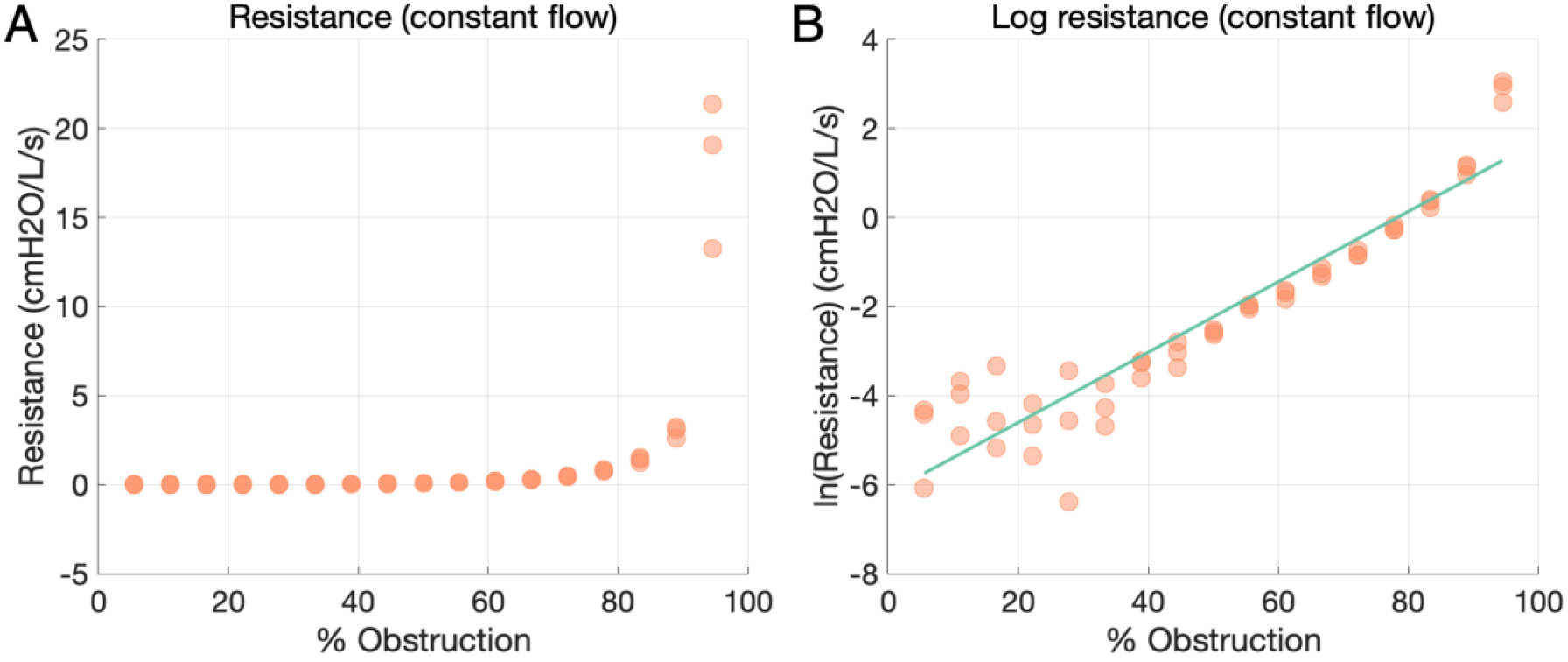
Inspiratory resistance as a function of obstruction under constant airflow. **A)** Effective resistance is plotted in terms of the percentage of airway obstruction. Resistance is seen to increase exponentially with obstruction from a value of about 70%. **B)** Log resistance as a function of airway obstruction. The magnitude of resistance is well fitted as a log-linear function of airway obstruction.

## Discussion

We present a novel psychophysical method and custom experimental apparatus for the streamlined and efficient estimation of respiroceptive sensitivity. To achieve this, we used accessible and low-cost 3D printing methods to develop an apparatus which can flexibly, safely, and reliably deliver inspiratory or expiratory loads entirely through a stimulus PC and sanitary circuit. By expanding well-established Bayesian psychometric approaches, we further developed a novel psychophysical task for estimating respiroceptive thresholds, the respiratory resistance sensitivity task (RRST). Our results show that the RRST provides highly reliable measurement of respiroceptive thresholds in just 20-30 minutes, and is minimally aversive for participants. Further, by enabling the rapid collection of many near-threshold trials in a 2IFC design, the RRST can flexibly dissociate interoceptive sensitivity, precision, and bias, and is well suited for model-based research in interoceptive perceptual decision-making over both perceptual and metacognitive dimensions (see e.g., (Allen et al., 2021; Allen, Levy, et al., 2019; Unal et al., 2021).

### The perception of breathing

A general issue in interoception research is the inability to dissociate sensitivity from response bias. This dissociation can be made by controlling the information present in the stimulus (i.e., the signal-to-noise ratio) in a precise way and quantifying the change in sensitivity resulting from increases in signal. However, the signal in most interoceptive sensory modalities, such as cardiac or gastric sensations, are not directly amenable to experimental control in the absence of highly invasive procedures such as distention of the esophagus, stomach and rectum (Jones et al., 2003; van Dyck et al., 2016; Yuan et al., 2003). This poses a fundamental limitation on studies of interoception, as well-established techniques for studying perception (e.g., psychophysics, signal detection theory) become difficult to apply in the absence of control over the physical signal. Further, human introspection is rife with cognitive and perceptual biases, meaning that purely subjective methods cannot identify the channel capacity or true sensitivity of the interoceptive system. While subjective biases themselves are of interest, this is a fundamental limitation for interoception research, where identifying the objective sensitivity to interoceptive stimuli may prove valuable both for unravelling the fundamental neural circuits underlying interoceptive perception, and for identifying maladaptive interoceptive sensitivity, which could be wholly orthogonal to interoceptive belief. As such, predominant approaches to measuring interoceptive perception either rely on difficult to interpret measures such as heart-beat counting (Brener & Ring, 2016; Schandry, 1981), or use highly invasive procedures such as gastric intubation (Stephan et al., 2003).

The respiratory domain presents interoception research both with unique opportunities and challenges. Respiratory sensation is itself a complex mix of interoceptive and exteroceptive sensory-motor and chemosensitive channels. During normal ventilation, air is mechanically pumped into the lungs by the distension and contraction of the diaphragm, which pushes air in and out of the lungs much like a blacksmith’s bellows. Throughout the inspiratory-expiratory cycle, the passage of air through the mouth, nose, and upper airway is communicated by thermosensory and tactile receptors, and the rhythmic expansion and contraction of the lungs is associated with the activation of stretch receptors located throughout the diaphragm, chest wall, and surrounding organs (Schroijen et al., 2020). This rhythmic information is communicated to the medullary and pontine brainstem nuclei, the somatosensory cortex, and higher-order structures such as the insula (Davenport & Vovk, 2009; Schroijen et al., 2020; von Leupoldt et al., 2008). Additionally, deviations in the concentrations of blood gases (carbon dioxide and oxygen) are communicated to the viscerosensory nuclei of the brain by chemosensory pathways, generating the interoceptive sensation of “air hunger” (Guz, 1997; Manning & Schwartzstein, 1995).

The complex physiological and neuroanatomical pathways subscribed by RI are therefore fundamentally multi-modal, and a mixture of interoceptive and exteroceptive sensations arising from the airway, blood, and skeletomuscular system. Alterations in either inspiratory or expiratory resistance can be expected to interact with this system at multiple levels: through the sensation of pressure at the lips, mouth, and upper airway, through the sensation of increased diaphragmatic effort, and in the case of sustained (i.e., non-discrete) loads, through the sensation of (likely) increased carbon dioxide in the bloodstream. A measure aiming at characterizing respiroception therefore needs to be flexible and fine-grained enough to resolve the sensations produced by these convergent breathing-related signals.

### Benefits of the RRST

A primary benefit of the RRST approach is that resistive stimuli can be flexibly manipulated to identify the minimal stimulus a participant can reliably discriminate. Ecologically speaking, the process of detecting respiratory loads operationalized by the RRST and similar procedures is not unlike that which accompanies a serious chest cold, where we might notice that our breathing has become more laboured due to the partial obstruction of the airway by mucous or inflammatory swelling. Respiratory resistance tasks therefore present an ecologically valid means by which to dissociate the objective sensitivity of airway monitoring.

However, as others have noted (Miller & Davenport, 2015), increasing the effort associated with respiration is for most individuals an inherently aversive process. This, coupled with the coarse granularity of most previous resistance-based tasks, meant that the estimation of resistive thresholds was a slow, painstaking process requiring many trials. By leveraging our unique apparatus, the RRST is able to adjust the intensity of the respiratory stimulus (e.g., the amount of airway resistance) in a highly granular manner. This opens the possibility to use Bayesian or other adaptive psychophysical techniques to optimize the delivered stimulus on each trial, greatly increasing the speed and precision with which respiratory psychophysical parameters such as threshold and precision can be estimated. Further, once threshold has been identified, the experimenter can flexibly adapt stimuli to each participant, delivering fine-grained control over task difficulty (i.e., error rates) and ensuring that task stimuli are equivalent across participants while controlling for variations in sensitivity (see **Supplementary Figure 4**). This, together with the 2IFC design of the RRST, is an important feature in particular for the estimation and modelling of respiratory metacognition where uncontrolled type 1 error rates can strongly bias estimates of metacognitive sensitivity and/or efficiency (Fleming & Lau, 2014; Guggenmos, 2021). Further, by maintaining fine-grained control over stimuli, the RRST ensures that excessively large resistive loads can be avoided and keeps total data acquisition times at just 20 - 30 minutes per participant, limiting the aversiveness of the task. This is especially important for clinical populations, such as persons with asthma or anxiety, where previous methods may simply be too challenging to apply in the very populations within whom RI abnormalities are likely to be of the greatest interest.

By use of a 2IFC design, our task is optimised for the measure of respiratory perceptual sensitivity. Single alternative (1IFC, or yes/no) tasks, in which a single stimulus is presented and the observer decides whether a signal was present or absent, are criterion-dependent. This reflects the observation that the participant needs to adopt a criterion, or a stimulus level at which they will begin to respond ‘yes’. Due to this dependency, 1IFC tasks characterise response bias as the propensity to respond that a signal is present, reflecting either an unconscious bias, or a response strategy (Morgan et al., 2012). In a 2IFC design such as that used in the RRST, this response bias manifests as a tendency to respond ‘first interval’ or ‘second interval’ more often. In both of these cases, the use of the signal detection theoretic d’ can allow for the dissociation of sensitivity from response bias.

The RRST brings several further practical benefits. The introduction of a visual pacing aid contributes to respiratory entrainment, assisting participants to pace their inhalations in a consistent manner. This likely contributes to the good same-day retest reliability we observed here (see **Figure 4D**), as well as to the relatively low levels of aversiveness reported by the participants. By encouraging quick, shallow and even breaths, most of the participants in the study were able to avoid hyperventilation and feelings of breathlessness, even following 30 minutes of testing. Finally, the components necessary in order to build the RRST apparatus are inexpensive and easily accessible through online vendors, and the shell can be 3D printed using standard commercial 3D printers, such as those available at the maker labs found at many Universities. We make all components, schematics and software available on GitHub, with the intention that others can build the device independently.

### Limitations

The task presented here evaluates inspiratory resistance sensitivity across cognitive levels, by measuring both type 1 perceptual performance (i.e., resistance sensitivity as measured by % airway obstruction at threshold, and psychometric function slope parameter) and type 2 metacognitive performance (measured by mRatio, or aROC). It is possible that variation in breathing patterns between participants results in substantial variations in airflow and therefore, in different values of pressure, at the same airway obstruction level (see **Supplementary Figure 3**). Here we introduced a number of features to mitigate the effect of variations in respiratory pattern. First, we designed a visual respiratory entrainment stimulus to standardise depth of inspirations between participants. Secondly, each trial of the task requires two short inhalations in quick succession, thereby regularizing respiratory frequency. Nevertheless, the variation in respiratory patterns, especially within clinical populations, leads us to recommend caution when interpreting type 1 results from the RRST, especially when the perceptual sensitivity is the primary variable of interest. In these cases, it would be recommended to record pressure and flow data during the task, to enable estimates of the psychometric parameters as a function of inspiratory pressure as a more nuanced measure, rather than of % airway obstruction, since it accounts for the inter- and intra-individual variability of air flow and likely reflects the effective stimulus intensity on each trial more accurately.

A further potential concern is the tolerance to the task, as prolonged exposure to inspiratory resistances was previously shown to be aversive to participants. Here, we regularly recorded subjective ratings of task aversiveness, as well as subjective ratings of breathlessness, dizziness and asthma symptoms at the end of each session. While a small subset of participants did find the task aversive, our results show that the RRST is well tolerated by most participants. Furthermore, individual levels of self-reported aversiveness were found to be uncorrelated with perceptual and metacognitive parameters, indicating that these parameters are orthogonal with respect to task aversiveness.

### Conclusion

Here we have presented the RRST, a novel method for measuring respiratory interoception-related factors across perceptual and metacognitive levels. By leveraging a custom-designed apparatus and Bayesian adaptive psychophysical algorithm, the RRST can reliably estimate threshold, slope and signal theoretic measures alongside metacognitive factors in under 30 minutes of testing. This short testing time combined with minimal aversiveness, opens up the possibility of studying respiratory interoception in clinical populations.

## Acknowledgements

NN, MB, NB, NL, and MA are supported by a Lundbeckfonden Fellowship (under Grant [R272-2017-4345]). MA is supported by the AIAS-COFUND II fellowship programme that is supported by the Marie Skłodowska-Curie actions under the European Union’s Horizon 2020 (under Grant [754513]), and the Aarhus University Research Foundation. OKH (née Faull) is supported by a Rutherford Discovery Research Fellowship awarded by the Royal Society of New Zealand, and the Department of Psychology at the University of Otago. FF is supported by a European Research Council grant (ERC-StG-948838).

## Supplementary Material

**Supplementary Figure 1:**
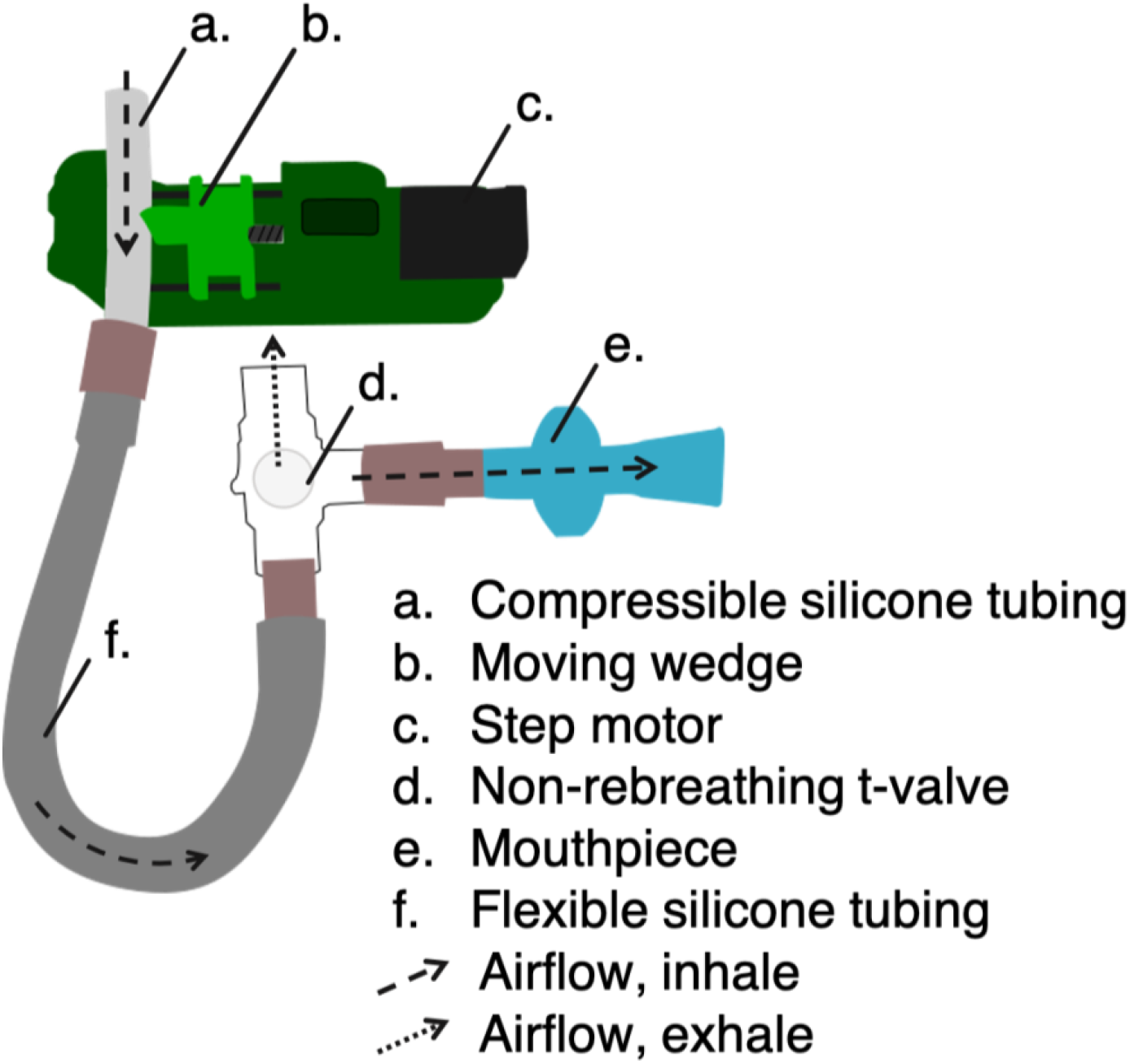
Detailed schematic illustrating the RRST apparatus. The apparatus (green) consists of a step motor (c) that turns a screw so as to move a wedge piece (b) back and forth. In this way, the wedge (b) can push on a piece of compressible silicone tubing (a), thereby obstructing it and effectively restricting the aperture. The breathing circuit consists of the silicone tubing (a), a flexible medical grade tube (f) and non-rebreathing 3-way T-valve. Participants inhale through the circuit using a single-use mouthpiece (e)

**Supplementary Figure 2.**
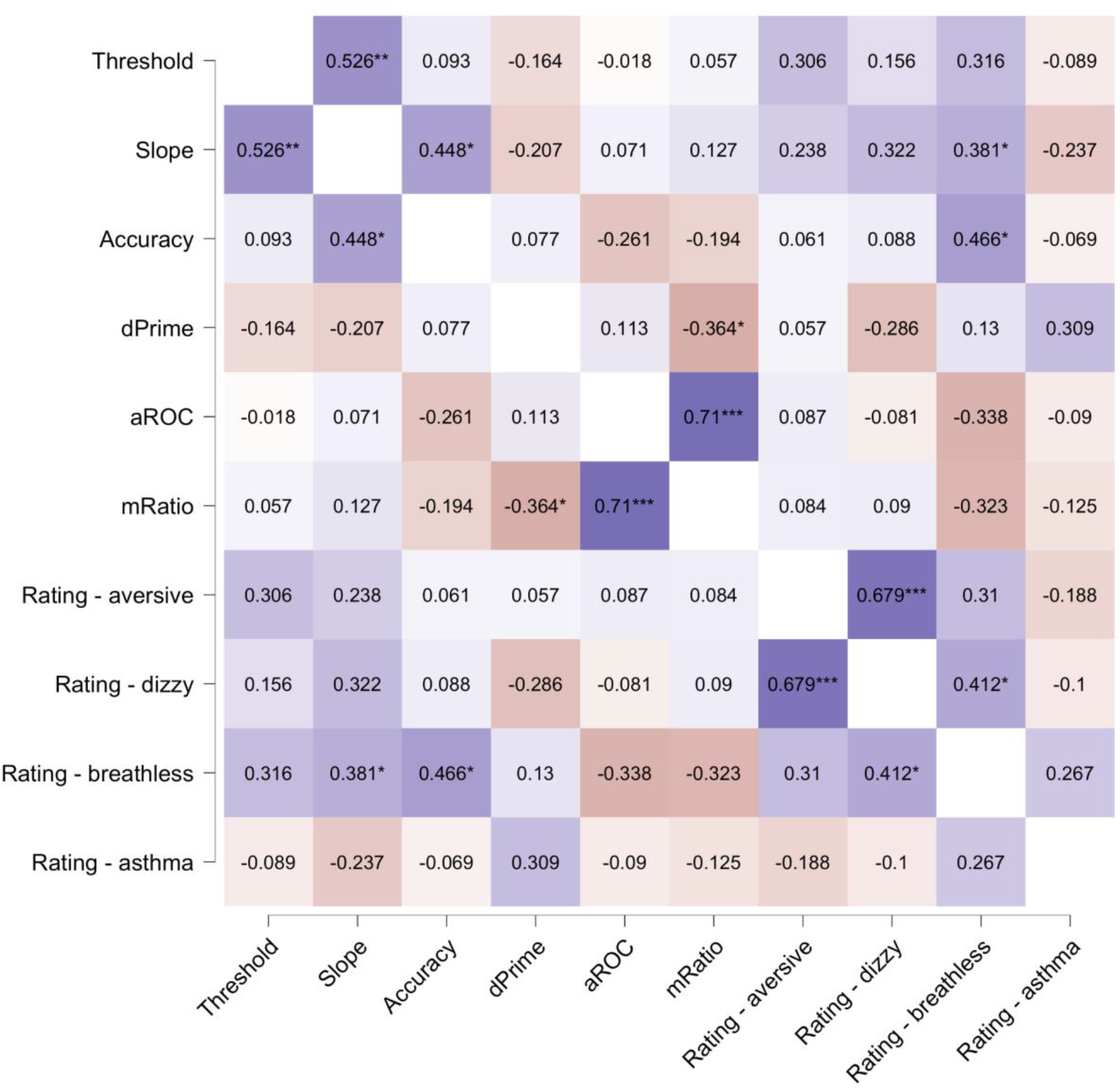
Correlation heatmap of task parameters and subjective ratings. We observed a correlation between threshold and slope estimates (ρ = 0.56), as well as between slope and task accuracy (ρ = 0.45). While threshold estimates were not related to subjective rating of aversiveness, dizziness or breathlessness, slope and accuracy were moderately related to ratings of breathlessness (ρ = 0.38 and ρ = 0.47, respectively). The metacognition variables aROC and mRatio correlate strongly (ρ = 0.71), as do ratings of dizziness and overall aversiveness on the task (ρ=0.68). Spearman’s rho correlations are represented. It should be noted, however, that our design is underpowered for this analyses and these results should therefor be treated as exploratory pending further replication in larger samples.

### Parts List

**Supplementary table 1:**
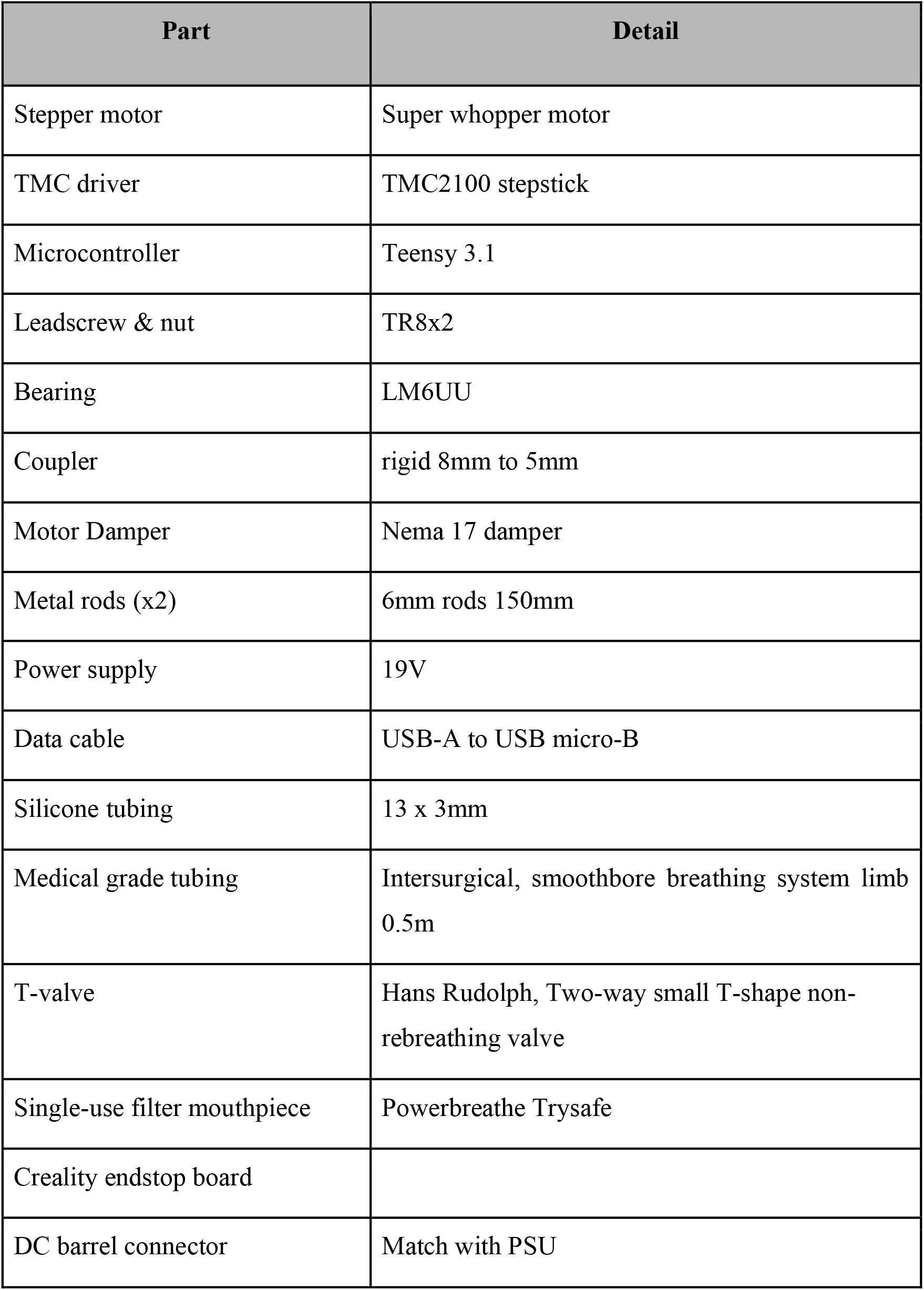
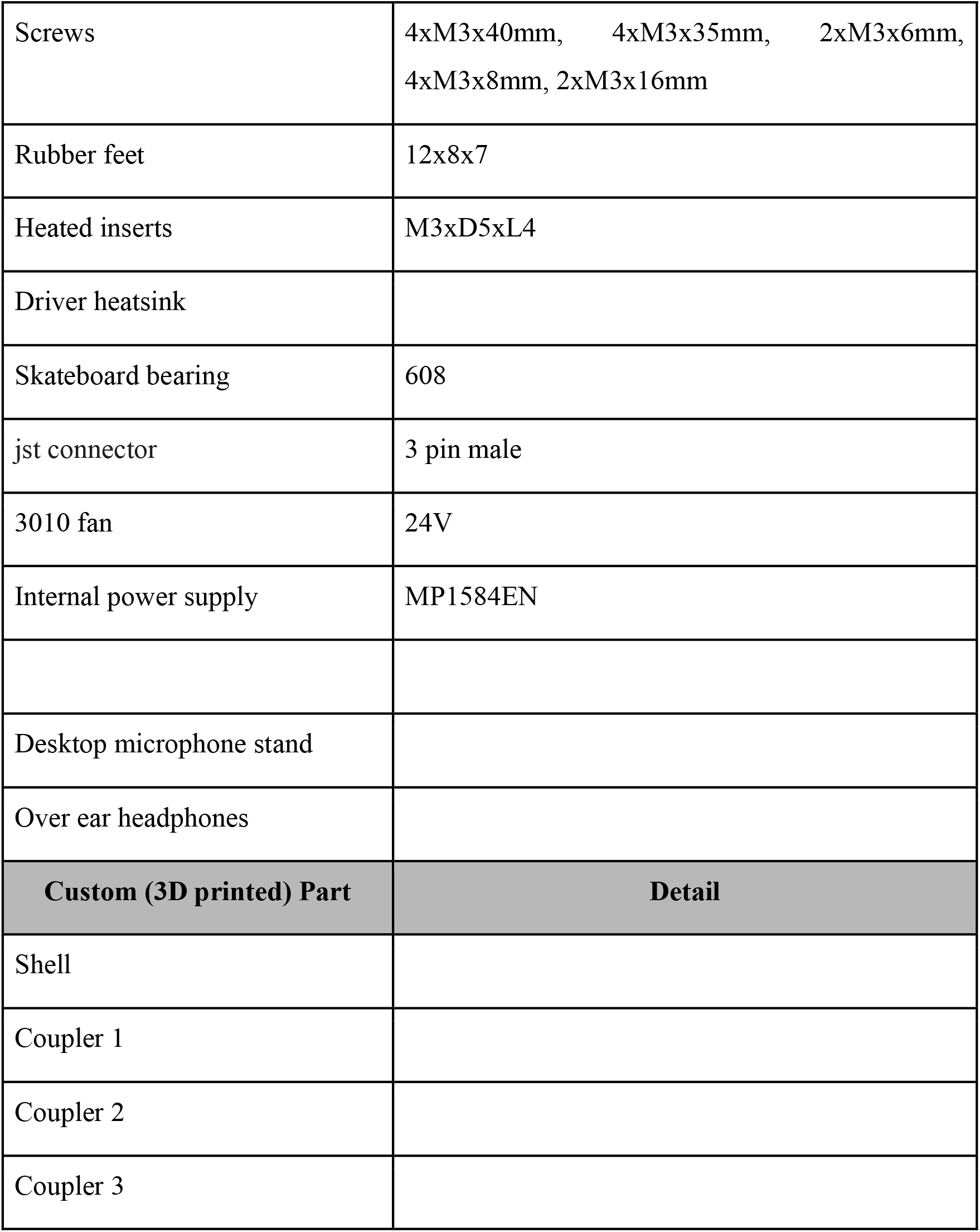
List of parts and components necessary for RRST apparatus.

### Task validation using physiological measures

The precise measurement of respiratory resistance thresholds is dependent on both reproducible resistive stimuli delivered by the apparatus (see **Figure 7**, main text), as well as reliable effective resistance values despite individual variability in breathing dynamics. To evaluate resistances produced by the RRST during real-life testing, we measured pressure and flow throughout sessions of the task.

#### Supplementary Methods

The data for this follow-up study was collected at a second site (University of Otago, New Zealand, Ethics number 20/CEN/168: Approval given by the New Zealand Health and Disability Ethics Committee). Fifteen participants (12 females, average age of 23.1 years ± 5.5 (SD)) completed a session of the RRST, while differential pressure and flow generated in the respiratory circuit were recorded (see main text Methods - Physiological measures for details).

#### Supplementary Results

Since we observed that the resistance produced on each step of airway obstruction is well fit by a log-linear function (Results - Inspiratory resistance), the results here are represented on a log scale as well. **Supplementary Figure 3** shows resistance (**A**) and pressure (**B**) measurements on individual trials for 15 sessions of the RRST completed by different participants. During natural performance of the task, the logarithm of both resistance and pressure is found to increase linearly with obstruction level.

**Supplementary Figure 3:**
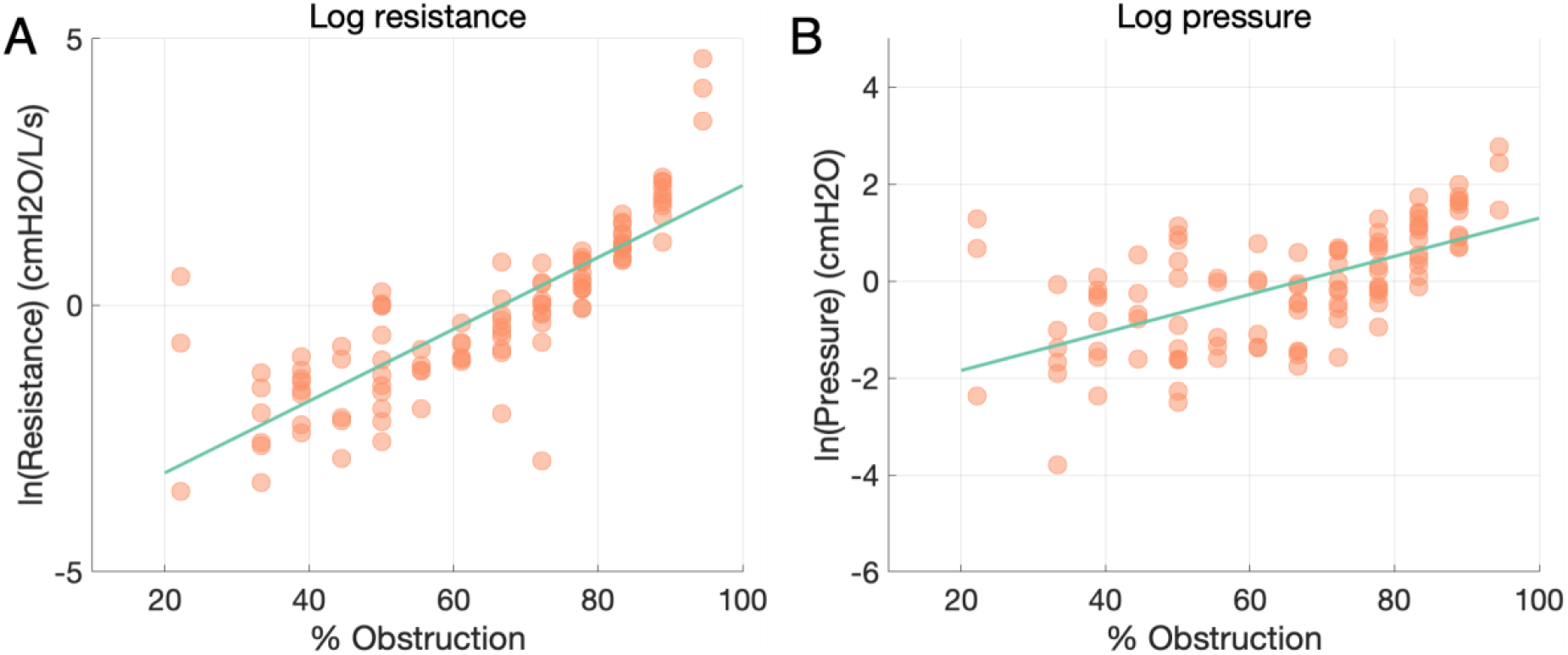
RRST resistance and pressure as a function of obstruction for participants. **A)** Natural log resistance (cm H_2_O/L.s^−1^) as a function of airway obstruction (r = 0.82, p < 0.0001). **B)** Natural log pressure (cm H_2_O) as a function of airway obstruction. Data points are participant means across trials at each level of obstruction (r = 0.62, p < 0.0001).

### Comparison to Filter Detection Task

We wanted to compare performance on the RRST to an existing measure of respiratory interoception, the Filter Detection Task (Harrison, Garfinkel, et al., 2021), to evaluate whether psychometric estimates (i.e., thresholds), metacognitive variables and accuracy relate between testing methods.

#### Methods

The data for this comparison was collected at a second site (University of Otago, New Zealand, see Supplementary material, Task validation using Physiological measures), in the same sample of participants as described in the previous section. Briefly, in the FDT between 0 and 12 spirometry filter head filters (each providing a resistance load of 0.42 cm H_2_O/L.s^−1^) are attached to a circuit, which the participant inhales through. The baseline resistance consisting of an empty/dummy filter was applied either in the first interval or second interval of three breaths, with the resistance load applied in the other interval of three breaths, for a total of six breaths in each trial. Calibration trials were performed before the task to determine the starting filter number at the participant’s perceptual threshold. A total of 60 trials were completed by each participant. On each trial, participants take two sets of three breaths on the circuit, and then decide whether the first or second set of breaths carried a greater resistance (i.e., a 2IFC task). The number of inspiratory resistance filters was determined using a staircase method implemented in MatLab, and was readjusted throughout the trials to maintain a task accuracy of 65 - 80%.

#### Supplementary Analyses

We first compared average physiological measures of pressure and resistance, as well as type 1 and type 2 performance variables between each task using paired-samples t-tests. We further conducted exploratory Pearson correlational analyses inter-relating FDT and RRST average resistance and pressure, as well as subjective confidence ratings and metacognition scores (a ROC). All analyses were Bonferroni corrected for multiple comparisons. For raw confidence analyses, RRST values were first divided by 10 to equate them to the 10 - point FDT scale. These analyses can be reproduced using the FDTvsRRST.jasp JASP data file found on the project GitHub: https://github.com/embodied-computation-group/RespiroceptionMethodsPaper/blob/main/suppAnalyses/FDTvsRRST.jasp

#### Results

Summary statistics describing physiological variables, accuracy, and metacognition are described below in Supplementary Table 2:

**Supplementary Table 2:** Means and Variances for key FDT and RRST Measures.

|  | FDT (SD) | RRST (SD) | t(13) | *p* | Cohen's *d* |
| --- | --- | --- | --- | --- | --- |
| Pressure (cm H<sub>2</sub>O) | 1.21 (13.16) | 2.75 (2.27) | -2.386 | 0.101 | -0.472 |
| Resistance (H<sub>2</sub>O/L.s<sup>-1</sup>) | 1.82 (1.21) | 7.92 (13.16) | -1.766 | 0.033 | -0.638 |
| Accuracy | 71% (9%) | 79% (1.5%) | -1.161 | 0.267 | -0.310 |
| Confidence | 5.2 (1.6) | 5.7 (1.4) | -3.196 | 0.007 | -0.854 |
| Metacognition (a ROC) | 0.63 (0.07) | 0.61 (0.05) | 0.759 | 0.462 | 0.203 |

*Note.* Student's *t*-test. FDT = Filter Detection Task, RRST = Respiratory Resistance Sensitivity Task.

In general, average performance parameters across the two tasks were highly similar, with the exception of increased average resistance and confidence in the RRST. No significant correlations were found between perceptual or metacognitive sensitivity, average resistance, or pressure on the two tasks (all ps > 0.05). We did observe a significant correlation between subjective confidence in the two tasks, Pearson’s *r(14)* = 0.75, p = 0.002.

While the RRST by design holds accuracy (hit rate) at 80% (± 1.5% SD), performance varies more widely in the FDT (mean accuracy 71% ± 9% SD), with 6 times higher standard deviation of accuracy on the FDT as compared to the RRST (**Supplementary Figure 4 A**). A tight control of accuracy is desirable for analyses interested in type 2 performance, as it allows for the estimation of metacognitive variables, independently of contamination by differences in type 1 performance (Guggenmos, 2021; Xue et al., 2021). Mean confidence ratings are found to correlate between the RRST and FDT (**Supplementary Figure 4 B**). This is in agreement with previous results showing that confidence ratings relate highly across modalities (e.g., Mazancieux et al., 2018).

**Supplementary Figure 4:**
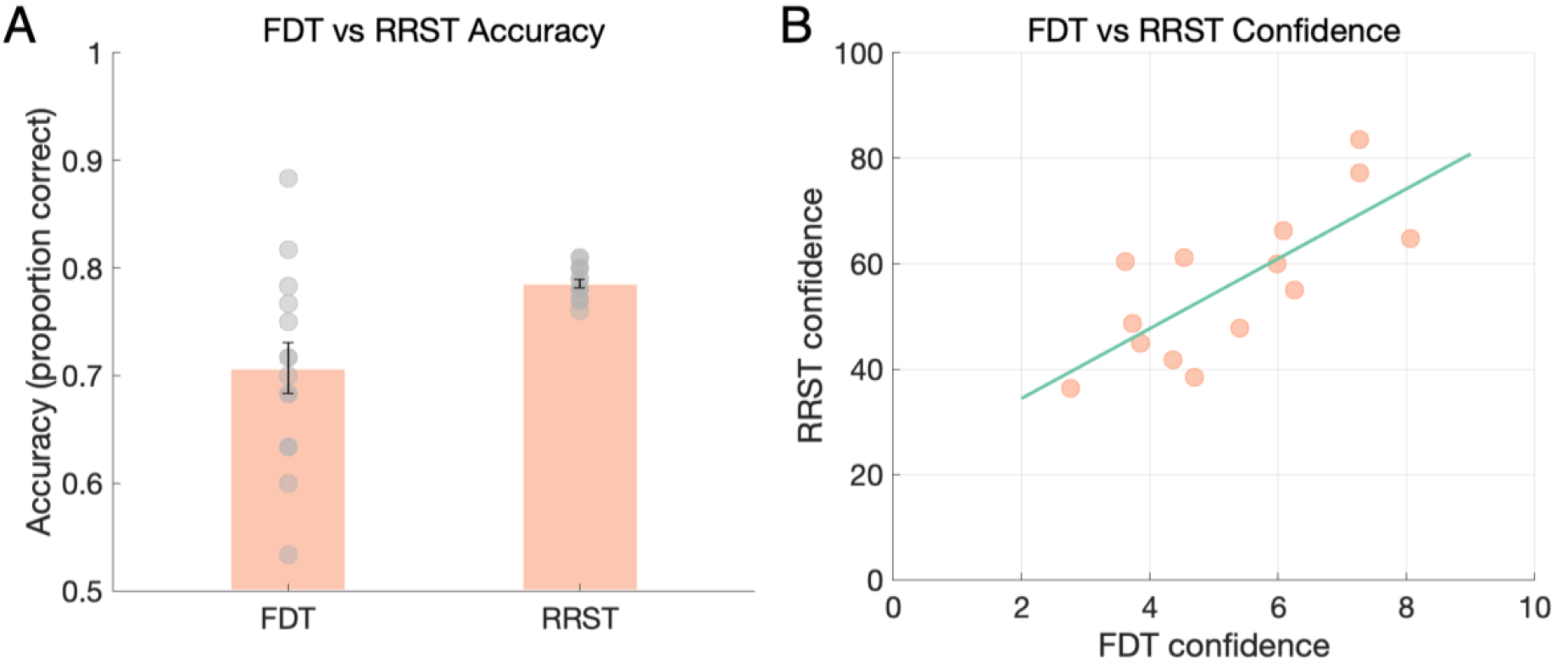
Comparison of FDT and RRST type 1 and type 2 variables. **A)** The RRST achieves a greater degree of control over accuracy, facilitating the comparison of metacognitive variables. Points denote overall task accuracies for each of 15 participants, bar height indicates mean accuracy across participants, and error bars indicate standard deviation. **B)** Confidence ratings correlate between FDT and RRST. Points represent mean confidence scores for each of 15 participants. Line denotes best fit of a linear regression (*R* = 0.75, *p* = 0.002).

Finally, for illustration purposes, we provide example respiratory physiological traces under constant flow and for an exemplary experimental participant, in Supplementary Figures 5 & 6, below.

**Supplementary Figure 5:**
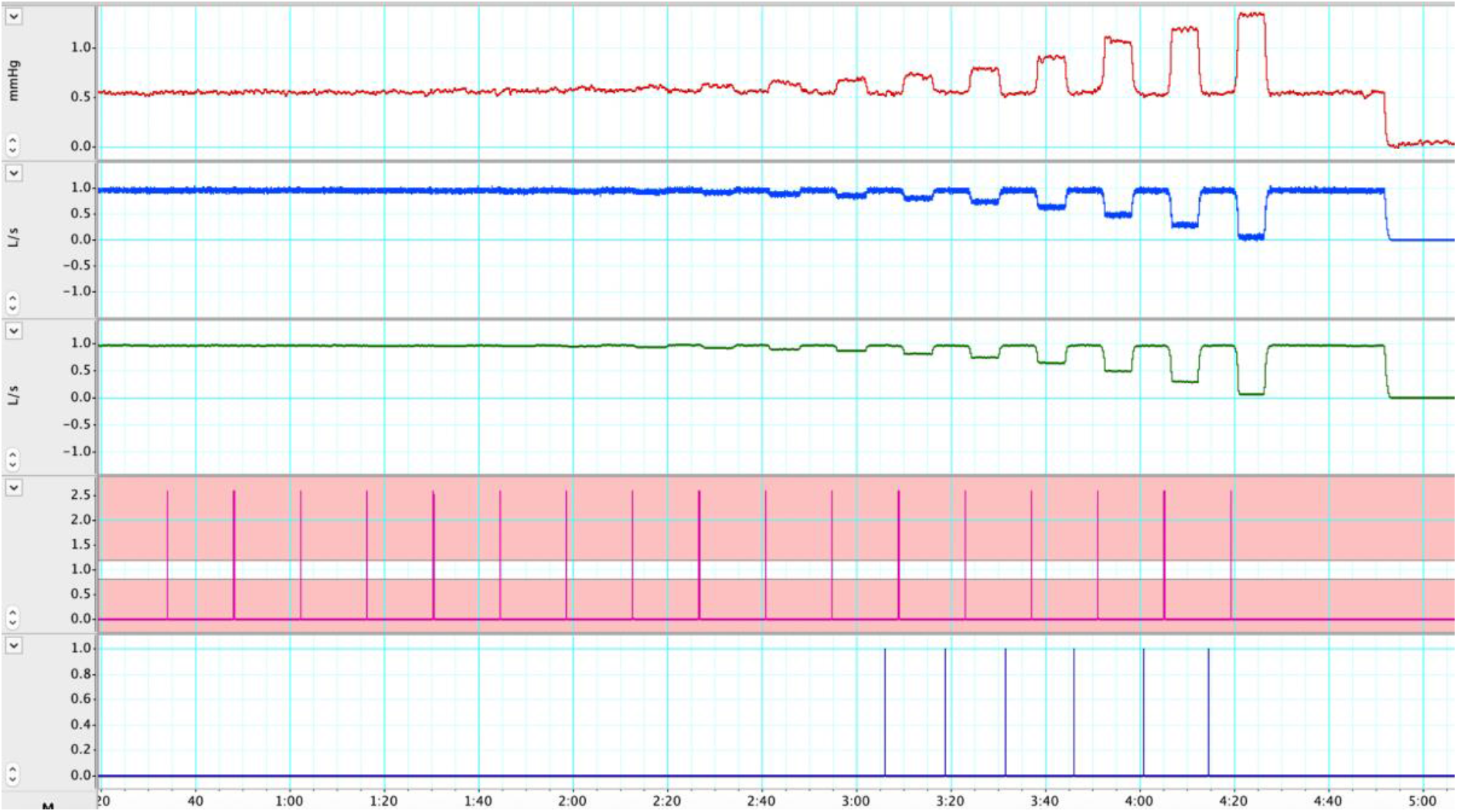
Constant flow mechanical measures RRST. Constant flow data during each step of the RRST. Pressure increases in log scale for each step. Top line = inspiratory pressure trace (mm Hg), second line = raw inspiratory flow trace (L/s), third line = smoothed inspiratory flow trace, fourth line = triggers indicating the onset of the inspiratory resistance stimuli, bottom line = triggers indicating an automatically-detected inspiration (adjustable depending on the analysis). Data collected in LabChart (version 8; ADInstruments).

**Supplementary Figure 6:**
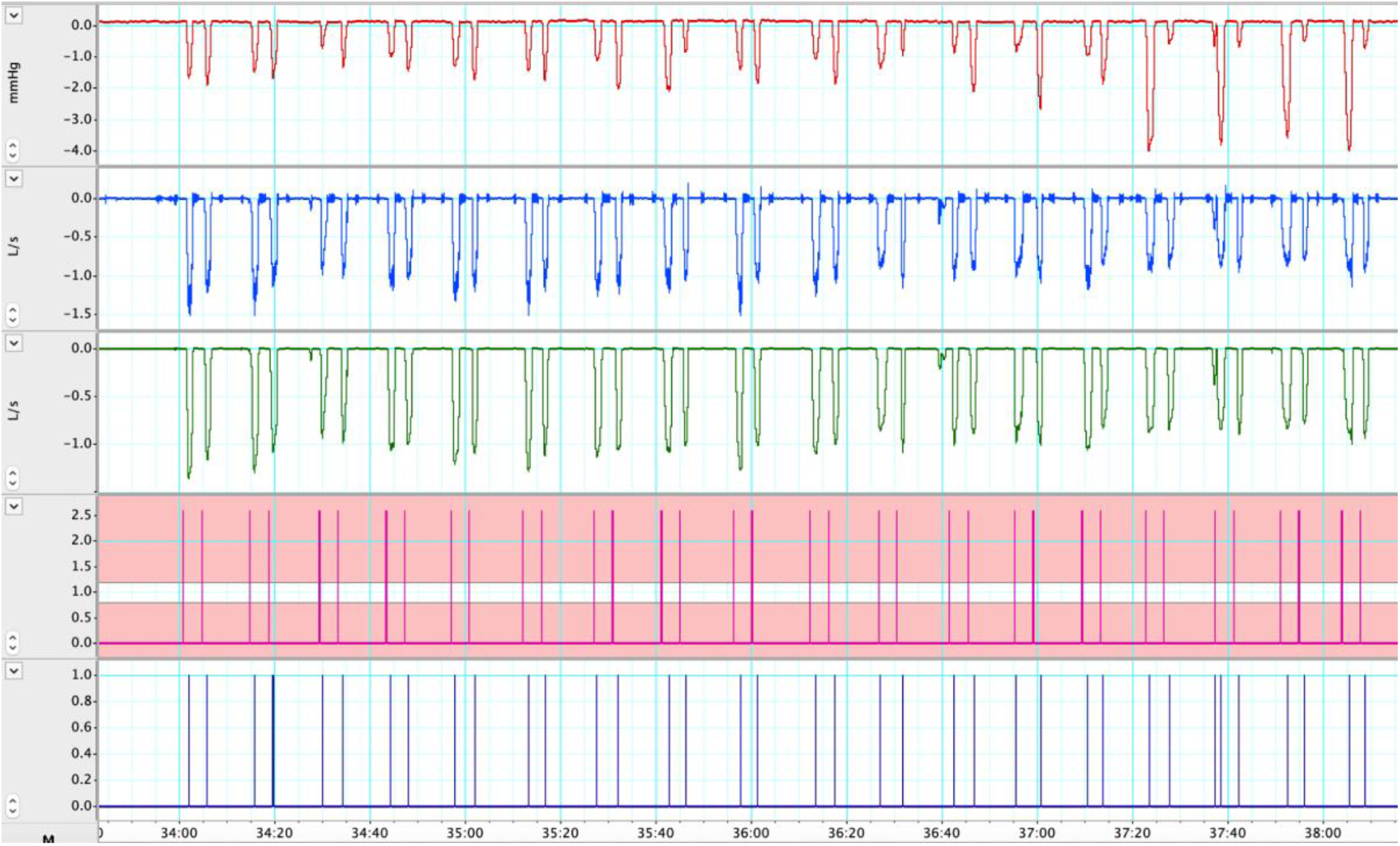
Example participant data RRST. Example participant data during the RRST. Top line = inspiratory pressure trace (mm Hg), second line = raw inspiratory flow trace (L/s), third line = smoothed inspiratory flow trace, fourth line = triggers indicating the onset of the inspiratory resistance stimuli, bottom line = triggers indicating an automatically-detected inspiration (adjustable depending on the analysis). Data collected in LabChart (version 8; ADInstruments).

## References

Allen, M. (2020). Unravelling the Neurobiology of Interoceptive Inference. Trends in Cognitive Sciences, 24(4), 265–266. https://doi.org/10.1016/j.tics.2020.02.002

Allen, M., Legrand, N., Correa, C. M. C., & Fardo, F. (2020). Thinking through prior bodies: Autonomic uncertainty and interoceptive self-inference. Behavioral and Brain Sciences, 43. https://doi.org/10.1017/S0140525X19002899

Allen, M., Levy, A., Parr, T., & Friston, K. J. (2019). In the Body’s Eye: The Computational Anatomy of Interoceptive Inference. BioRxiv, 603928. https://doi.org/10.1101/603928

Allen, M., Poggiali, D., Whitaker, K., Marshall, T. R., & Kievit, R. A. (2019). Raincloud plots: A multi-platform tool for robust data visualization. Wellcome Open Research, 4, 63. https://doi.org/10.12688/wellcomeopenres.15191.1

Allen, M., Varga, S., & Heck, D. H. (2021). Respiratory Rhythms of the Predictive Mind. PsyArXiv. https://doi.org/10.31234/osf.io/38bpw

Bailey, P. H. (2004). The Dyspnea-Anxiety-Dyspnea Cycle—COPD Patients’ Stories of Breathlessness: “It’s Scary /When you Can’t Breathe”. Qualitative Health Research, 14(6), 760–778. https://doi.org/10.1177/1049732304265973

Bennett, E. D., Jayson, M. I., Rubenstein, D., & Campbell, E. J. (1962). The ability of man to detect added non-elastic loads to breathing. Clinical Science, 23, 155–162.

Bogaerts, K., Millen, A., Li, W., De Peuter, S., Van Diest, I., Vlemincx, E., Fannes, S., & Van den Bergh, O. (2008). High symptom reporters are less interoceptively accurate in a symptom-related context. Journal of Psychosomatic Research, 65(5), 417–424. https://doi.org/10.1016/j.jpsychores.2008.03.019

Brainard, D. H. (1997). The Psychophysics Toolbox. Spatial Vision, 10(4), 433–436. https://doi.org/10.1163/156856897X00357

Brener, J., & Ring, C. (2016). Towards a psychophysics of interoceptive processes: The measurement of heartbeat detection. Philosophical Transactions of the Royal Society B: Biological Sciences, 371(1708), 20160015. https://doi.org/10.1098/rstb.2016.0015

Critchley, H. D., & Garfinkel, S. N. (2017). Interoception and emotion. Current Opinion in Psychology, 17, 7–14. https://doi.org/10.1016/j.copsyc.2017.04.020

Dahme, B., Richter, R., & Mass, R. (1996). Interoception of respiratory resistance in asthmatic patients. Biological Psychology, 42(1), 215–229. https://doi.org/10.1016/0301-0511(95)05156-2

Davenport, P. W., & Vovk, A. (2009). Cortical and subcortical central neural pathways in respiratory sensations. Respiratory Physiology & Neurobiology, 167(1), 72–86. https://doi.org/10.1016/j.resp.2008.10.001

Fleming, S. M. (2017). HMeta-d: Hierarchical Bayesian estimation of metacognitive efficiency from confidence ratings. Neuroscience of Consciousness, 2017(1). https://doi.org/10.1093/nc/nix007

Fleming, S. M., & Lau, H. C. (2014). How to measure metacognition. Frontiers in Human Neuroscience, 8. https://doi.org/10.3389/fnhum.2014.00443

Garfinkel, S. N., Manassei, M. F., Hamilton-Fletcher, G., In den Bosch, Y., Critchley, H. D., & Engels, M. (2016). Interoceptive dimensions across cardiac and respiratory axes. Philosophical Transactions of the Royal Society B: Biological Sciences, 371(1708), 20160014.

Guz, A. (1997). Brain, breathing and breathlessness. Respiration Physiology, 109(3), 197–204. https://doi.org/10.1016/S0034-5687(97)00050-9

Harrison, O. K., Garfinkel, S. N., Marlow, L., Finnegan, S. L., Marino, S., Köchli, L., Allen, M., Finnemann, J., Keur-Huizinga, L., Harrison, S. J., Stephan, K. E., Pattinson, K. T. S., & Fleming, S. M. (2021). The Filter Detection Task for measurement of breathing-related interoception and metacognition. Biological Psychology, 165, 108185. https://doi.org/10.1016/j.biopsycho.2021.108185

Harrison, O. K., Garfinkel, S. N., Marlow, L., Finnegan, S., Marino, S., Nanz, L., Allen, M., Finnemann, J., Keur-Huizinga, L., Harrison, S. J., Stephan, K. E., Pattinson, K., & Fleming, S. M. (2020). The Filter Detection Task for measurement of breathing-related interoception and metacognition. BioRxiv, 2020.06.29.176941.https://doi.org/10.1101/2020.06.29.176941

Harrison, O. K., Marlow, L., Finnegan, S., Ainsworth, B., & Pattinson, K. T. S. (2020). Dissociating breathlessness symptoms from mood in asthma (p. 2020.07.15.204289). https://doi.org/10.1101/2020.07.15.204289

Harrison, O. K., Nanz, L., Marino, S., Lüchinger, R., Hennel, F., Hess, A. J., Frässle, S., Iglesias, S., Vinckier, F., Petzschner, F., Harrison, S. J., & Stephan, K. E. (2021). Interoception of breathing and its relationship with anxiety. BioRxiv, 2021.03.24.436881. https://doi.org/10.1101/2021.03.24.436881

Harver, A., Katkin, E. S., & Bloch, E. (1993). Signal-detection outcomes on heartbeat and respiratory resistance detection tasks in male and female subjects. Psychophysiology, 30(3), 223–230. https://doi.org/10.1111/j.1469-8986.1993.tb03347.x

James, W. (1884). What is an Emotion? Mind, 9(34), 188–205. JSTOR.

James, W. (1894). Discussion: The physical basis of emotion. Psychological Review, 1(5), 516–529. https://doi.org/10.1037/h0065078

Jones, M. P., Hoffman, S., Shah, D., Patel, K., & Ebert, C. C. (2003). The water load test: Observations from healthy controls and patients with functional dyspepsia. American Journal of Physiology-Gastrointestinal and Liver Physiology, 284(6), G896–G904. https://doi.org/10.1152/ajpgi.00361.2002

Khalsa, S. S., Adolphs, R., Cameron, O. G., Critchley, H. D., Davenport, P. W., Feinstein, J. S., Feusner, J. D., Garfinkel, S. N., Lane, R. D., Mehling, W. E., Meuret, A. E., Nemeroff, C. B., Oppenheimer, S., Petzschner, F. H., Pollatos, O., Rhudy, J. L., Schramm, L. P., Simmons, W. K., Stein, M. B., … Paulus, M. P. (2018). Interoception and Mental Health: A Roadmap. Biological Psychiatry. Cognitive Neuroscience and Neuroimaging, 3(6), 501–513. https://doi.org/10.1016/j.bpsc.2017.12.004

Khalsa, S. S., & Lapidus, R. C. (2016). Can Interoception Improve the Pragmatic Search for Biomarkers in Psychiatry? Frontiers in Psychiatry, 7. https://doi.org/10.3389/fpsyt.2016.00121

King-Smith, P. E., Grigsby, S. S., Vingrys, A. J., Benes, S. C., & Supowit, A. (1994). Efficient and unbiased modifications of the QUEST threshold method: Theory, simulations, experimental evaluation and practical implementation. Vision Research, 34(7), 885–912. https://doi.org/10.1016/0042-6989(94)90039-6

Kleiner, M., Brainard, D., Pelli, D., Ingling, A., Murray, R., & Broussard, C. (2007). What’s new in psychtoolbox-3. Perception, 36(14), 1–16.

Kontsevich, L. L., & Tyler, C. W. (1999). Bayesian adaptive estimation of psychometric slope and threshold. Vision Research, 39(16), 2729–2737. https://doi.org/10.1016/S0042-6989(98)00285-5

Maniscalco, B., & Lau, H. (2012). A signal detection theoretic approach for estimating metacognitive sensitivity from confidence ratings. Consciousness and Cognition, 21(1), 422–430. https://doi.org/10.1016/j.concog.2011.09.021

Manning, H. L., & Schwartzstein, R. M. (1995). Pathophysiology of Dyspnea. New England Journal of Medicine, 333 (23), 1547–1553. https://doi.org/10.1056/NEJM199512073332307

Mazancieux, A., Fleming, S. M., Souchay, C., & Moulin, C. (2018). Retrospective confidence judgments across tasks: Domain-general processes underlying metacognitive accuracy [Preprint]. PsyArXiv. https://doi.org/10.31234/osf.io/dr7ba

Meuret, A. E., Rosenfield, D., Hofmann, S. G., Suvak, M. K., & Roth, W. T. (2009). Changes in respiration mediate changes in fear of bodily sensations in panic disorder. Journal of Psychiatric Research, 43(6), 634–641. https://doi.org/10.1016/j.jpsychires.2008.08.003

Morélot-Panzini, C., Demoule, A., Straus, C., Zelter, M., Derenne, J.-P., Willer, J.-C., & Similowski, T. (2007). Dyspnea as a Noxious Sensation: Inspiratory Threshold Loading May Trigger Diffuse Noxious Inhibitory Controls in Humans. Journal of Neurophysiology, 97(2), 1396–1404. https://doi.org/10.1152/jn.00116.2006

Morgan, M., Dillenburger, B., Raphael, S., & Solomon, J. A. (2012). Observers can voluntarily shift their psychometric functions without losing sensitivity. Attention, Perception & Psychophysics, 74(1), 185–193. https://doi.org/10.3758/s13414-011-0222-7

Nikolova, N., Waade, P. T., Friston, K. J., & Allen, M. (2021). What Might Interoceptive Inference Reveal about Consciousness? Review of Philosophy and Psychology. https://doi.org/10.1007/s13164-021-00580-3

Owens, A. P., Allen, M., Ondobaka, S., & Friston, K. J. (2018). Interoceptive inference: From computational neuroscience to clinic. Neuroscience & Biobehavioral Reviews, 90, 174–183. https://doi.org/10.1016/j.neubiorev.2018.04.017

Pelli, D. G. (1997). The VideoToolbox software for visual psychophysics: Transforming numbers into movies. Spatial Vision, 10(4), 437–442. https://doi.org/10.1163/156856897X00366

Prins, N., & Kingdom, F. A. A. (2018). Applying the Model-Comparison Approach to Test Specific Research Hypotheses in Psychophysical Research Using the Palamedes Toolbox. Frontiers in Psychology, 9. https://doi.org/10.3389/fpsyg.2018.01250

Schandry, R. (1981). Heart Beat Perception and Emotional Experience. Psychophysiology, 18(4), 483–488. https://doi.org/10.1111/j.1469-8986.1981.tb02486.x

Schroijen, M., Davenport, P. W., Bergh, O. V.den, & Diest, I. V. (2020). The Sensation of Breathing. In Cotes’ Lung Function (pp. 407–422). John Wiley & Sons, Ltd. https://doi.org/10.1002/9781118597309.ch22

Sherrington, C. (1952). The Integrative Action of the Nervous System. CUP Archive. https://books.google.dk/books?hl=en&lr=&id=CRU8AAAAIAAJ&oi=fnd&pg=PR7&dq=Sherrington,+C.+(1952).+The+integrative+action+of+the+nervous+system&ots=KBRtrndr41&sig=zQmk7Bfcw364FEPZYmaNZezrvEQ&redir_esc=y#v=onepage&q=Sherrington%2C%20C.%20(1952).%20The%20integrative%20action%20of%20the%20nervous%20system&f=false

Stephan, E., Pardo, J. V., Faris, P. L., Hartman, B. K., Kim, S. W., Ivanov, E. H., Daughters, R. S., Costello, P. A., & Goodale, R. L. (2003). Functional neuroimaging of gastric distention. Journal of Gastrointestinal Surgery, 7(6), 740–749. https://doi.org/10.1016/S1091-255X(03)00071-4

Tiller, J., Pain, M., & Biddle, N. (1987). Anxiety Disorder and Perception of Inspiratory Resistive Loads. Chest, 91(4), 547–551. https://doi.org/10.1378/chest.91.4.547

Tweeddale, P. M., Rowbottom, I., & McHardy, G. J. R. (1994). Breathing retraining: Effect on anxiety and depression scores in behavioural breathlessness. Journal of Psychosomatic Research, 38(1), 11–21. https://doi.org/10.1016/0022-3999(94)90004-3

Unal, O., Eren, O. C., Alkan, G., Petzschner, F. H., Yao, Y., & Stephan, K. E. (2021). Inference on homeostatic belief precision. Biological Psychology, 165, 108190. https://doi.org/10.1016/j.biopsycho.2021.108190

Vaitl, D. (1996). Interoception. Biological Psychology, 42 (1), 1–27. https://doi.org/10.1016/0301-0511(95)05144-9

van Dyck, Z., Schulz, A., Blechert, J., Herbert, B. M., Lutz, A. P. C., & Vögele, C. (2021). Gastric interoception and gastric myoelectrical activity in bulimia nervosa and binge-eating disorder. International Journal of Eating Disorders, 54(7), 1106–1115. https://doi.org/10.1002/eat.23291

van Dyck, Z., Vögele, C., Blechert, J., Lutz, A. P. C., Schulz, A., & Herbert, B. M. (2016). The Water Load Test As a Measure of Gastric Interoception: Development of a Two-Stage Protocol and Application to a Healthy Female Population. PLoS ONE, 11(9). https://doi.org/10.1371/journal.pone.0163574

von Leupoldt, A., Sommer, T., Kegat, S., Baumann, H. J., Klose, H., Dahme, B., & Büchel, C. (2008). The Unpleasantness of Perceived Dyspnea Is Processed in the Anterior Insula and Amygdala. American Journal of Respiratory and Critical Care Medicine, 177(9), 1026–1032. https://doi.org/10.1164/rccm.200712-1821OC

Watson, A. B., & Pelli, D. G. (1983). Quest: A Bayesian adaptive psychometric method. Perception & Psychophysics, 33(2, 113–120. https://doi.org/10.3758/BF03202828

Wiley, R. L., & Zechman, F. W. (1966). Perception of added airflow resistance in humans. Respiration Physiology, 2(1), 73–87. https://doi.org/10.1016/0034-5687(66)90039-9

Yuan, Y.-Z., Tao, R.-J., Xu, B., Sun, J., Chen, K.-M., Miao, F., Zhang, Z.-W., & Xu, J.-Y. (2003). Functional brain imaging in irritable bowel syndrome with rectal balloon-distention by using fMRI. World Journal of Gastroenterology: WJG, 9(6), 1356–1360. https://doi.org/10.3748/wjg.v9.i6.1356

